# Bidirectional cooperation between Ubtf1 and SL1 determines RNA Polymerase I promoter recognition *in cell* and is negatively affected in the UBTF-E210K neuroregression syndrome

**DOI:** 10.1101/2021.06.07.447350

**Authors:** Michel G. Tremblay, Dany S. Sibai, Melissa Valère, Jean-Clément Mars, Frédéric Lessard, Roderick T. Hori, Mohammad M. Khan, Victor Y. Stefanovsky, Mark S. Ledoux, Tom Moss

## Abstract

Transcription of the ∼200 mouse and human ribosomal RNA genes (rDNA) by RNA Polymerase I (RPI/PolR1) accounts for 80% of total cellular RNA, around 35% of all nuclear RNA synthesis, and determines the cytoplasmic ribosome complement. It is therefore a major factor controlling cell growth and its misfunction has been implicated in hypertrophic and developmental disorders. Activation of each rDNA repeat requires nucleosome replacement by the architectural multi-HMGbox factor UBTF to create a 15kbp nucleosome free region (NFR). Formation of this NFR is also essential for recruitment of the TBP-TAF_I_ factor SL1 and for preinitiation complex (PIC) formation at the gene and enhancer-associated promoters of the rDNA. However, these promoters show little sequence commonality and neither UBTF nor SL1 display significant DNA sequence binding specificity, making what drives PIC formation a mystery. Here we show that cooperation between SL1 and the longer UBTF1 splice variant generates the specificity required for rDNA promoter recognition *in cell*. We find that conditional deletion of the Taf1b subunit of SL1 causes a striking depletion UBTF at both rDNA promoters but not elsewhere across the rDNA. We also find that while both UBTF1 and −2 variants bind throughout the rDNA NFR, only UBTF1 is present with SL1 at the promoters. The data strongly suggest an induced-fit model of RPI promoter recognition in which UBTF1 plays an architectural role. Interestingly, a recurrent UBTF-E210K mutation and the cause of a pediatric neurodegeneration syndrome provides indirect support for this model. E210K knock-in cells show enhanced levels of the UBTF1 splice variant and a concomitant increase in active rDNA copies. In contrast, they also display reduced rDNA transcription and promoter recruitment of SL1. We suggest the underlying cause of the UBTF-E210K syndrome is therefore a reduction in cooperative UBTF1-SL1 promoter recruitment that may be partially compensated by enhanced rDNA activation.

## INTRODUCTION

The ribosomal RNA (rRNA) genes encode the catalytic and structural RNAs of the ribosome as a single 47S precursor. As such, transcription of these genes is a major determinant of cell growth, cell cycle progression and cell survival and an essential factor in the formation of hypertrophic diseases such as cancer (1). Dysregulation of rRNA genes is also the cause of a large range of developmental and neurological disorders that are often also associated with cancer (2–6). To develop treatment strategies for these diseases it is important that we command an understanding of how these genes are transcribed and regulated. Transcription of the several hundred tandemly repeated and essentially identical rRNA genes, the rDNA, is undertaken exclusively by RNA Polymerase I (RPI, Pol1, PolR1) and a set of basal transcription factors dedicated to this task. This strict correspondence of gene and polymerase has resulted in the rapid coevolution of rDNA promoters with basal factors, leading to a high degree of species specificity of the RPI transcription machinery (7–9). The functional uniqueness of the RPI machinery provides an obvious target for novel therapeutic approaches (10). However, what directs the RPI transcription machinery exclusively to the rDNA and how it is specifically recruited to both the major 47S pre-rRNA promoter and enhancer element despite these having little or no DNA sequence commonality are still not understood. Here we show that despite having little or no inherent DNA sequence selectivity, the multi-HMGbox Upstream Binding Factor (UBF/UBTF) plays a crucial role in targeting RPI preinitiation complex formation to the rDNA promoters *in vivo*. Further, we show that this UBTF function is compromised by an E210K mutation recently linked to a recurrent human pediatric neuroregression syndrome (6, 11–13).

The basal factors of the mammalian RPI transcription machinery include Selectivity Factor 1 (SL1), the multi-HMGbox factor UBTF/UBF and the RPI-associated initiation factor RRN3 (8, 14, 15) (Figure 1A). SL1 consists of the TATA-box Binding Protein (TBP) and the RPI specific TBP Associated Factors, TAFIA to D. Evolutionary variability of these TAFs determine their species-specific functions and that of RPI transcription, and for example human or mouse SL1 complexes are not functionally exchangeable (16–18). In contrast, UBTF is a highly conserved essential factor that is functionally exchangeable between human, mouse and to some extent even Xenopus (18–20). Our present understanding of how SL1 and UBTF function in rDNA transcription derives predominantly from cell-free studies. These have suggested sequential binding scenarios for the formation of the RPI preinitiation complex, whereby one or two dimers of UBTF bind across the rDNA promoter to provide a landing site for SL1, though the converse has also been suggested (21, 22). UBTF was found to bend and loop DNA, suggesting that it could bring together the two distal Upstream Promoter Element (UPE, aka UCE) and Core Elements of the RPI promoter to form such a landing site for SL1 (23). However, UBTF of itself does not display any significant degree of sequence specific binding, making its role in targeting SL1 difficult to understand (9). Indeed, UBTF binds continuously throughout the transcribed regions of the active rDNA genes, where it creates a 20 kbp long Nucleosome-Free Region (NFR) bounded upstream by CTCF and the enhancer associated Spacer Promoter, flanked by the nucleosomal InterGenic Spacer (IGS) (24, 25) (Figure 1B). Despite this, targeted gene inactivation has unequivocally shown Ubtf to be essential for recruitment of SL1 to both the major 47S promoter and the enhancer associated Spacer Promoter in mouse (24, 26).

**Figure 1.**
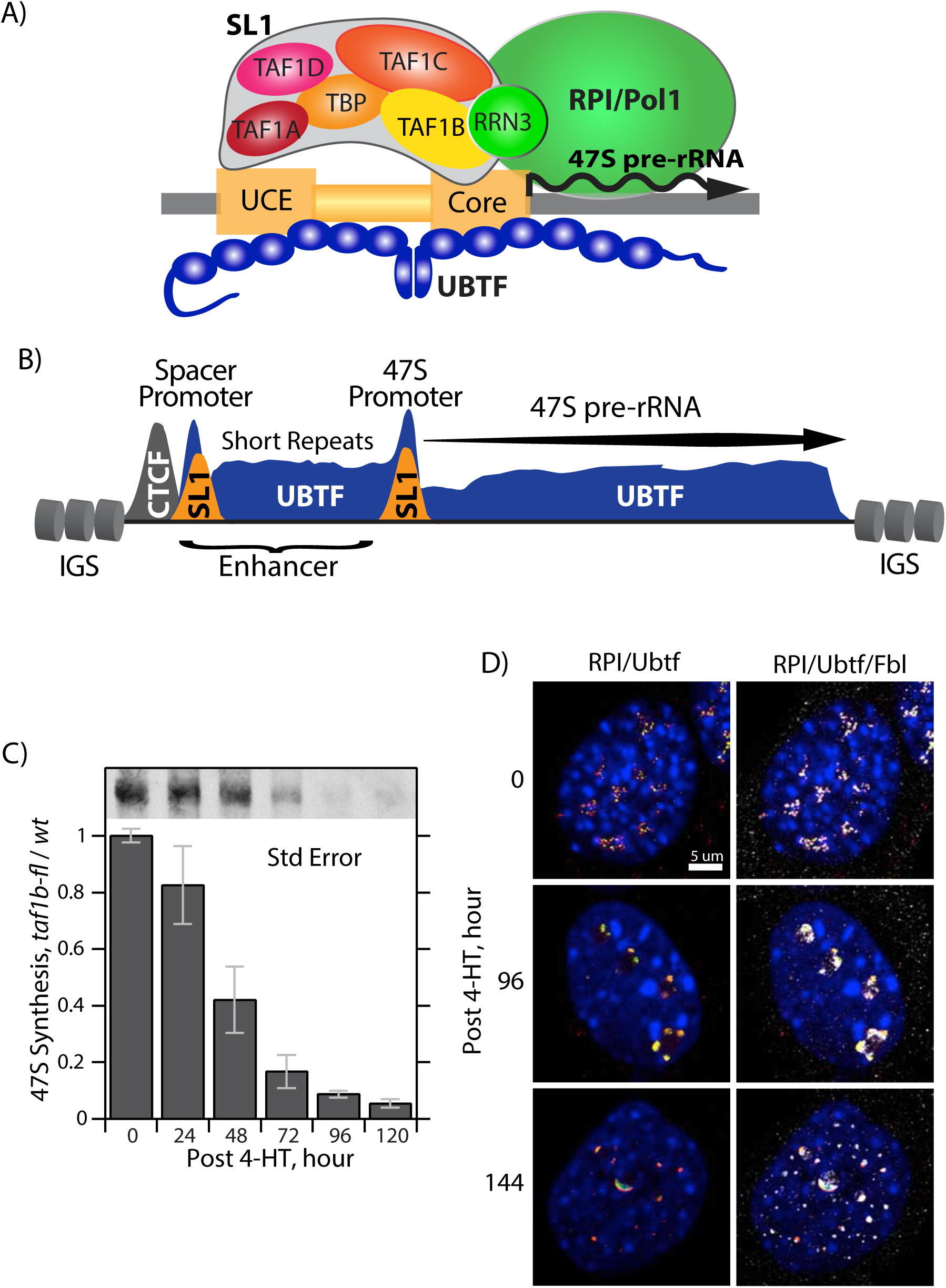
Genetic inactivation of the *taf1b* gene disrupts rRNA synthesis and nucleolar structure in conditional MEFs. A) Schematic of the basal factors of the RNA Polymerase I (RPI/Pol1) initiation complex and their assembly on the rDNA promoter. The diagram is not intended to indicate precise factor positioning. B) General organisation of the rDNA chromatin structure in mouse. The positions of Spacer and 47S Promoter duplications, the distribution of Ubtf, CTCF binding at the upstream boundary and the flanking nucleosomal IGS are indicated. C) Synthesis of 47S pre-rRNA in *taf1b* conditional MEFs. 47S pre-rRNA synthesis was determined by [3H]-uridine RNA metabolic labelling following homozygous *taf1b* deletion by 4-HT induction of ER-Cre. The upper inset panel shows a typical example of 47S pre-rRNA labelling in *taf1b^fl/fl^/p53^-/-^/ERcre^+/+^* MEFs at different times after ER-Cre induction. The lower histogram shows 47S labelling in *taf1b^fl/fl^/p53^-/-^/ERcre^+/+^* MEFs normalized to labelling in control *taf1b^wt/wt^/p53^-/-^/ERcre^+/+^* MEFs. The data shown in the histogram were derived from 6 independent analyses of MEFs derived from two floxed and two wild type embryos. Error bars indicate the SEM. See Figure S2 for the time course of *taf1b* deletion and Taf1b protein depletion. D) RPI, Ubtf and fibrillarin (FBL) indirect immunofluorescence labelling in *taf1b^fl/fl^/p53^-/-^/ERcre^+/+^* MEFs after 4-HT induction of *taf1b* deletion, see Figure S3 for a more detail.

Here we extend recent studies of Ubtf and Rrn3 (24, 26) to a conditional cell culture model for the Taf1b (TAF68) subunit of SL1, to the roles of the Ubtf variants and to a UBTF-E210K mutation recently identified as the cause of a recurrent pediatric neurodegeneration syndrome. The resulting data provide the first *in cell* test of the requirement for an RPI-specific TAF and a significant new insight into RPI preinitiation complex formation. The data resolve key questions surrounding rDNA promoter recognition and RPI pre-initiation complex formation by showing that the Ubtf1 splice variant displays a striking specificity for the RPI promoter sequences only when in the presence of a functional SL1. They further suggest an “induced-fit” model of promoter recognition in which UBTF plays an architectural role to model rDNA conformation to fit SL1 and hence catalyze its recruitment. Our findings further suggest that the fundamental cause of the UBTF-E210K pediatric neuroregression syndrome is a partial defect in SL1-Ubtf cooperation leading to reduced RPI preinitiation complex formation.

## RESULTS

### The Taf1B subunit of SL1 is essential for mouse development and rDNA transcription

Mouse lines carrying a targeted “Knockout First” insertion in the gene for Taf1B (Taf68), were established and these crossed to generate lines carrying either conditional *taf1b^flox^* or *taf1b*^Δ^ null-alleles (Figure S1). Mice heterozygous for a *taf1b*^Δ^ allele were found to be both viable and fertile and the null-allele was propagated at near Mendelian frequency (Table S1). However, no *Taf1b*^Δ/Δ^ homozygous offspring (pups) were identified and genotyping of embryos detected no *Taf1b*^Δ/Δ^ homozygotes at stages 6.5 and later. It was therefore concluded that *Taf1b* was essential for mouse development beyond blastula, see Supplementary data for more detail.

To determine the cellular effects of Taf1B loss, *taf1b-flox* mice (Figure S1A and B) were also crossed to introduce *ER-Cre* and *p53-null* alleles and conditional (*taf1b^fl/fl^/ER-Cre^+/+^/p53^-/-^*) and isogenic control (*taf1b^wt/wt^/ER-Cre^+/+^/p53^-/-^*) mouse embryonic fibroblasts (MEFs) were isolated as previously described (24, 27). Addition of 50 nM 4-hydroxytamoxifen (4-HT) to the conditional MEFs induced homozygous recombination of *Taf1b^flox^* alleles. This treatment strongly depleted the Taf1B protein already 24 h post 4-HT and essentially eliminated it 96 h post 4-HT. As expected for an integral subunit of SL1, depletion of Taf1B was paralleled by the strong suppression of 47S pre-rRNA synthesis (Figure 1C and Figure S2).

### Taf1B deletion induces nucleolar stress

Deletion of the *Taf1b* gene also led to a disruption of nucleolar structure characteristic of nucleolar stress. Prior to Taf1B depletion, immunofluorescence imaging (IF) of conditional MEFs revealed the typical punctate sub-nucleolar pattern of RPI and Ubtf overlapped by fibrillarin (FBL) staining indicative of transcriptionally active rDNA gene units (0h post 4-HT, in Figures 1D and S3). During Taf1B depletion, RPI and Ubtf staining collapsed into common intense foci that were often arranged in pairs around more central FBL staining (e.g. 96 and 120h post 4-HT in Figures 1D and S3). At later times, Ubtf became highly condensed and partially segregated from RPI while FBL dispersed throughout the nucleus (120 h and 144 h post 4-HT in Figures 1D and S3). These changes were consistent with the nucleolar changes previously observed on inactivation of RPI transcription either by Rrn3 gene deletion or CX-5461 drug inhibition (24, 28).

### Loss of Taf1B prevents SL1 recruitment but has only a small effect on “active” rDNA chromatin

Chromatin Immunoprecipitation (ChIP-qPCR) revealed that deletion of the Taf1b gene essentially eliminated Taf1b binding at the 47S and Spacer rDNA promoters in both MEFs and ESCs. It also prevented recruitment of the SL1 subunits TBP and Taf1c at both promoters (Figure 2B and S4). Thus, loss of Taf1b functionally inactivated SL1 and prevented preinitiation complex formation, explaining the suppression of rDNA transcription. In contrast, loss of Taf1b had only a limited effect on Ubtf binding across the 47S gene body where it predominantly replaces histone-based chromatin (24, 25) (see ETS and 28S amplicons in Figures 2B and S4). Consistent with this, psoralen accessibility crosslinking (PAC) indicated only a small reduction in the “active” form of rDNA chromatin that was previously shown to be dependent on Ubtf (24–26) (Figure 2C). Nevertheless, an increase in the mobility of the “active” PAC band on Taf1B loss suggested a higher degree of rDNA chromatin compaction, most probably related to the concomitant loss of RPI loading. A similar observation was made when RRN3 was inactivated (24, 29).

**Figure 2.**
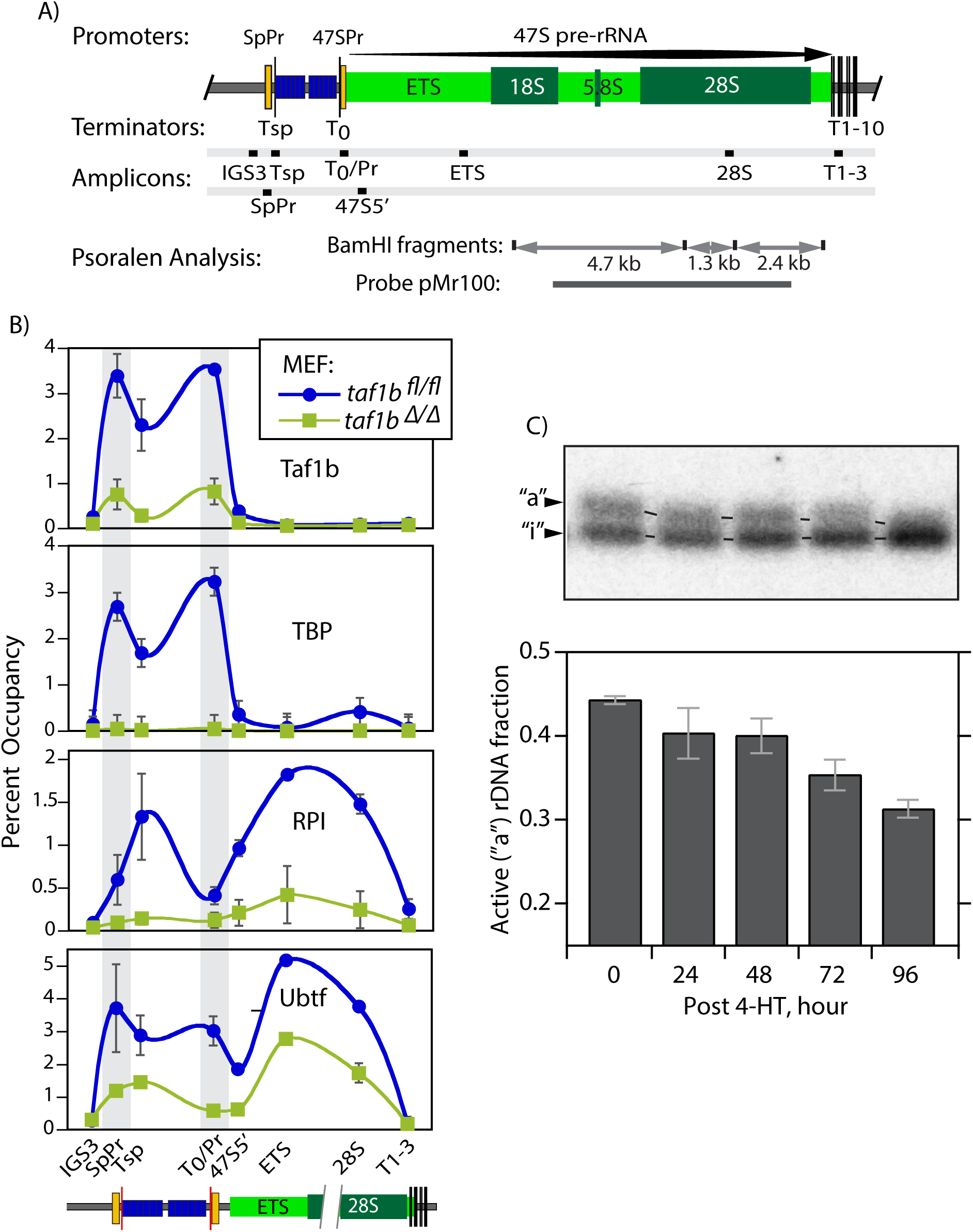
Loss of Taf1b abrogates RPI but not Ubtf recruitment to the rDNA. A) Organisation of the mouse rDNA locus indicating the positions of the Spacer (SpPr) and 47S (47SPr) promoter sequences, the RPI termination sites Tsp, T_0_ and T1-10, the extent 47S pre-rRNA coding region (light green) and the encoded 18, 5.8 and 28S rRNAs. The positions of the qPCR amplicons used in ChIP analyses are indicated, as are the rDNA fragments and pMr100 probe used in psoralen accessibility cross linking (PAC). B) ChIP-qPCR analysis of Taf1b, RPI and Ubtf occupancy at sites across the rDNA of *taf1b^fl/fl^/p53^-/-^/ERcre^+/+^* MEFs before and 5 days after *taf1b* inactivation by 4-HT treatment. The data derive from 3 biological ChIP replicas each analyzed by qPCR in triplicate. Figure S4 shows a similar ChIP analysis in *taf1b^fl/fl^/ERcre^+/+^* mESCs with mapping of Taf1b, Taf1c and TBP subunits of SL1. C) PAC reveals a gradual reduction in the amount and mobility of active rDNA chromatin following *taf1b* inactivation as in B. Upper panel shows a typical psoralen time course analysis of rDNA chromatin showing the lower mobility of the active “a” and higher mobility of the inactive “i” 1.3kbp BamHI-BamHI fragment from the rDNA 47S coding region. The lower histogram panel shows the mean active rDNA fraction estimated from curve fit analysis of the 1.3, 2.4 and 4.7kbp BamHI-BamHI rDNA fragment profiles in two biological replicas. Error bars indicate the SEM.

### The recruitment of Ubtf and SL1 to the rDNA promoters is highly cooperative

Though Taf1b depletion and the loss of SL1 recruitment to the rDNA had only a small effect on Ubtf binding across the gene body, inspection of the ChIP-qPCR mapping suggested that selective depletion did occur at both rDNA promoters (compare SpPr and T0/Pr with ETS and 28S amplicons in Figures 2B and S4). This was confirmed using the higher resolution of Deconvolution ChIP-Seq mapping (DChIP-Seq) (9, 24). Before Taf1b depletion, DChIP-Seq revealed overlapping peaks of SL1 (Taf1b) and Ubtf at both rDNA promoters in conditional MEFs (Figure 3A). Subsequent Taf1b depletion essentially eliminated it from the promoters but also strongly suppressed the overlapping peak of Ubtf, and this same effect was also seen in Taf1b conditional mESCs (Figure S5A). Despite the Taf1b-dependent loss of Ubtf from the rDNA promoters, its binding profile elsewhere across the rDNA was unaffected, though a generalized 25 to 50% loss of Ubtf respectively in mESCs and MEFs was observed. The dependence of Ubtf binding on Taf1b, and hence on functional SL1, was most evident in DChIP-Seq difference maps, which showed strong suppression of Ubtf specifically at 47S and Spacer promoters in both MEFs and mESCs types (Figure 3A, B and S5A, B).

**Figure 3.**
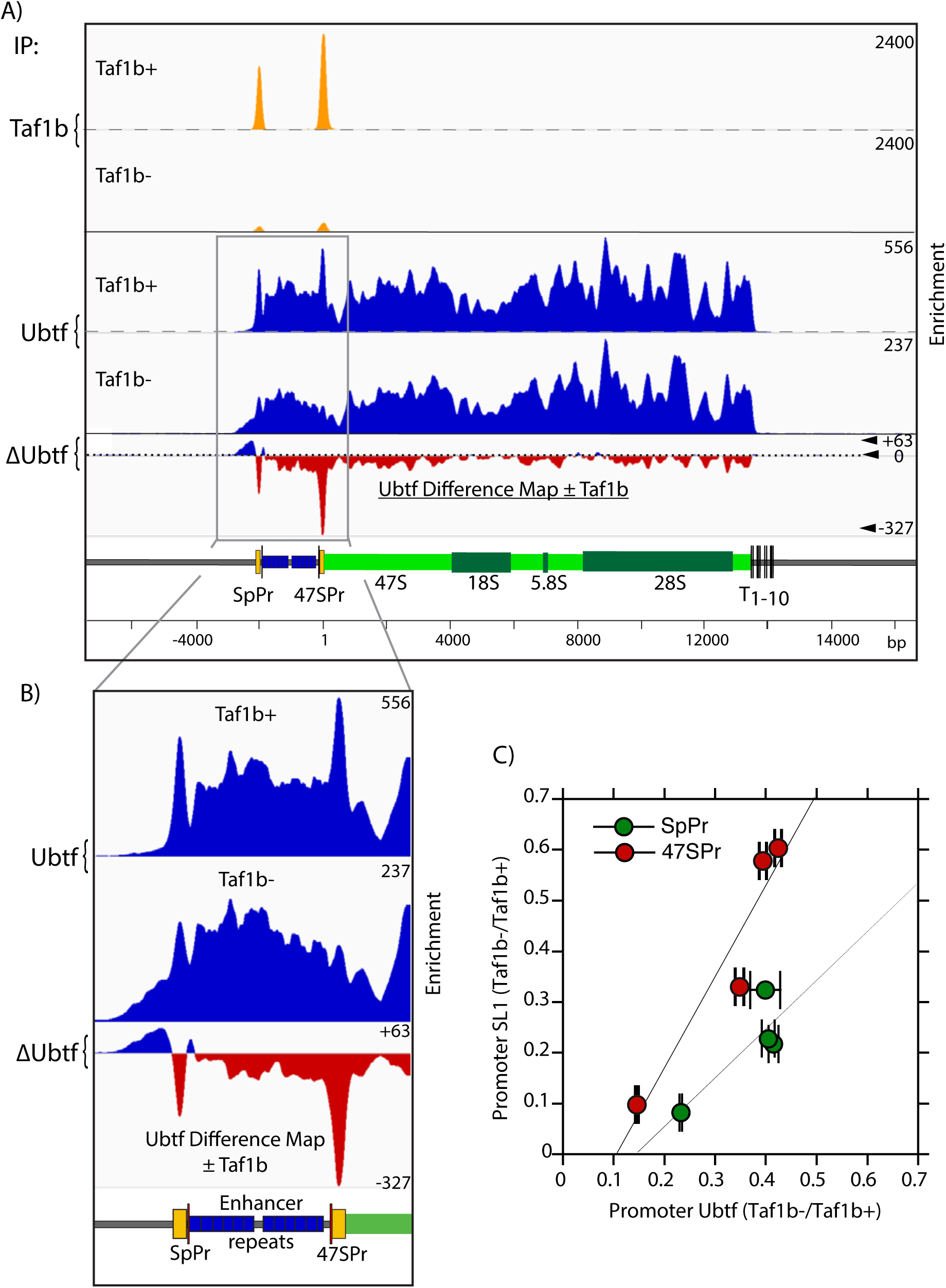
Taf1b loss induces depletion of Ubtf from both Spacer and 47S promoters but not from the adjacent enhancer repeats nor from the 47S gene body. A) DChIP-Seq analysis of Taf1b and Ubtf occupancy across the rDNA repeat in *taf1b^fl/fl^/p53^-/-^/ERcre^+/+^* MEFs before (Taf1b+) and 5 days after *taf1b* inactivation (Taf1b-). ΔUbtf indicates the difference map of Ubtf occupancy after Taf1b depletion minus the occupancy before Taf1b depletion. B) Magnified view of the DChIP mapping in A showing detail over the promoter and enhancer regions. C) Analysis of Spacer and 47S promoter occupancies reveals a direct proportionality between Taf1b and Ubtf. Four independent DChIP Ubtf and Taf1b data sets displaying different levels of Taf1b depletion were quantitatively analyzed for Ubtf and Taf1b promoter occupancy by peak fit, examples of which are shown in Figure S6. The fractions of SL1 (Taf1b) and Ubtf on each promoter after Taf1b depletion are plotted one against the other and reveal near linear relationships. Error bars in C show the SEM associated with peak fitting and.

The generation of DChIP-Seq profiles at differing degrees of Taf1b depletion allowed a quantitative estimate of the interdependence of Ubtf and SL1 binding at both rDNA promoters (see Materials and Methods and Figure S6). The data revealed near linear relationships between SL1 and Ubtf occupancy and confirmed that their binding was strongly interdependent at either promoter (Figure 3C). This was particularly striking given that these promoters share only 26% base sequence identity, no more than expected for two sequences chosen at random. Interestingly, the data (Figure 3C) also suggested a 2-fold difference in the relative SL1 : Ubtf stoichiometries between the Spacer and the 47S promoter, potentially a factor in their differential promoter strengths and functionalities.

### Only the longer of the two Ubtf variants is recruited to the rDNA promoters

The observation that Ubtf recruitment depended on SL1 specifically at the rDNA promoters but not elsewhere across the rDNA repeat suggested that the Ubtf variants might be important in this specificity. Mammals express two splice variants of Ubtf, both Ubtf1 and Ubtf2 encompass six tandem HMGbox DNA binding domain homologies but differ in HMGB-box2, a central segment of which is deleted in Ubtf2 (Figure 4A). While MEFs express both forms of Ubtf, ESCs naturally express exclusively Ubtf1 (Figure S5C). Promoter recruitment of Ubtf1 in these cells was found to be strongly suppressed on depletion functional SL1 (Figure S5A and B). However, this left open the question of whether or not promoter recruitment of Ubtf2 might also depended on SL1. To answer this question, we determined the distribution of each Ubtf variant across the rDNA in MEFs.

**Figure 4.**
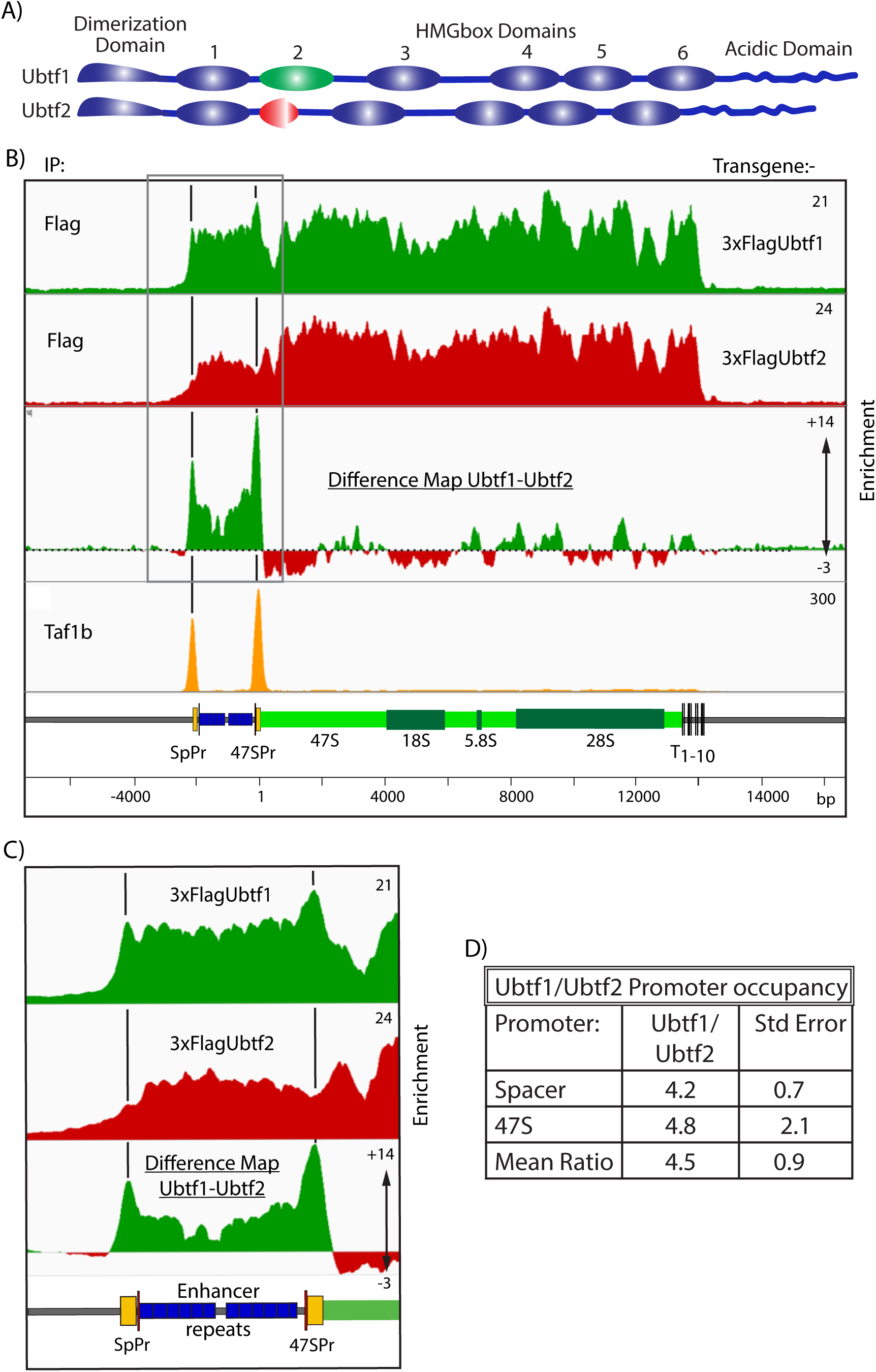
The rDNA promoters specifically recruit the Ubtf1 variant. A) Schematic representation of the domain structure of the Ubtf splice variants in mouse and human indicating the N-terminal dimerization, 6 HMGbox domains and the C-terminal Acidic Domain. B) DChIP-Seq mapping profiles of exogenously expressed 3xFLAG-Ubtf1 or Ubtf2, and endogenous Taf1b in NIH3T3 MEFs, see also Figure S7. The difference map of Ubtf1-Ubtf2 occupancies reveals a strong selectivity for Ubtf1 mapping precisely over the Spacer and 47S promoters. C) Magnified view of the DChIP mapping in B showing detail over the promoter and enhancer regions. D) Peak fit analysis of Ubtf1 and Ubtf2 occupancies over the Spacer and 47S promoter revealed that Ubtf1 was at least 4 times more prevalent at either promoter. The data derive from two biological replicas and the SEM is shown.

Pools of NIH3T3 MEF clones expressing 3xFLAG-tagged Ubtf1 or Ubtf2 at sub-endogenous levels were selected and subjected to DChIP-Seq mapping (Figure 4B and S7A and B). The profiles of the 3xFLAG-Ubtf1 and -Ubtf2 binding closely followed that of total endogenous Ubtf across most of the rDNA, however, it was significantly different at the Spacer and 47S promoters. Characteristic peaks of Ubtf were present at both promoters in the Ubtf1 profile but were absent in the Ubtf2 profile. This differential promoter binding was most evident in Ubtf1-Ubtf2 difference maps (Figure 4B and C, and S7B). Quantitative analysis of the variant Ubtf occupancy profiles (Experimental Procedures and Figure S8) showed greater than 4 times more Ubtf1 than Ubtf2 at the rDNA promoters (Figure 4D). However, our previous studies of the Ubtf-DNA complex show that a Ubtf dimer contacts contiguously 130 to 140 bp of DNA, arguing that each 150-170bp rDNA promoter could interact with at most two dimers of Ubtf (9, 23). Hence, the rDNA promoters must predominantly, if not exclusively, recruit Ubtf1. Further, since Ubtf1 was the sole variant present in ESCs and its promoter recruitment depended strongly on functional SL1 (Figure S5), it was concluded that promoter recognition and preinitiation complex formation *in vivo* specifically required the Ubtf1 variant.

The combined data showed that formation of the RPI preinitiation complex *in cell* involves a cooperation between Ubtf1 and SL1, and most surprisingly, this same cooperation occurred at both the Spacer and 47S promoters despite their unrelated base sequences. Since the only difference between Ubtf1 and Ubtf2 lies in the structure of HMGbox2, this domain must play a key role in Ubtf-SL1 cooperation and RPI promoter recognition.

### An HMGbox2 mutation linked to neuroregression potentially affects Ubtf interactions

An E>K mutation at residue 210 in HMGbox2 of Ubtf was recently shown to be the cause of a recurrent human pediatric neuroregression syndrome (6, 11–13). The key role of HMGbox2 revealed by our study suggested that this mutation might affect the formation of the RPI preinitiation complex *in vivo* and possibly explain the origin of this syndrome. Unfortunately, as yet the structure of HMGbox2 has not been determined experimentally. However, despite a high degree of primary sequence variability, HMGboxes display very similar tertiary structures and DNA contacts, making them accessible to molecular modelling (summarized in Figure S9A). Modelling of Ubtf-HMGbox2 revealed a typical HMGB saddle structure with basic residues K198, 200 and 211 lining the DNA binding underside (Figure S9B). Significantly, the sidechain of residue K211, a highly conserved minor groove contact in other HMGboxes, was predicted to be correctly oriented towards the DNA. In contrast, the sidechain of the immediately adjacent E210 residue was predicted to point away from the DNA and lay on the seat of the HMGbox saddle. Furthermore, this predicted sidechain position was unaffected by the E210K mutation (Figure S9C). We concluded that the E210K mutation was extremely unlikely to affect HMGbox2 interactions with the DNA. However, the mutation would create a significant change in the electrostatic surface potential of the seat of HMGbox2 (Figure S9D), suggesting that it could well affect interactions with other factors such as SL1.

### The UBTF HMGbox2 E210K mutation suppresses 47S rRNA synthesis in a MEF model

Given that the sequences of human and mouse UBTF are 99% identical, we took advantage of a recently generated *Ubtf^E210K^* mouse knock-in model. Mice homozygous for the E210K mutation are viable but exhibit behavioral abnormalities that worsen with increasing postnatal age (details will be described elsewhere). *Ubtf^E210K/E210K^* MEFs were isolated from these mice and found to proliferate somewhat more slowly than MEFs from isogenic wild type littermates, doubling times of 35h and 31h respectively (Figure 5A). Metabolic RNA labelling also revealed a >40% lower rate of *de novo* 47S pre-rRNA synthesis in the mutant as compared to the wild type MEFs (Figure 5B), however, no overt rRNA processing defects were detected (Figure S10A). The mutant MEFs also contained 30% less total cellular RNA, (∼80% of which is of course rRNA), than wild type MEFs (Figure 5C). Thus, the E210K mutation in Ubtf significantly reduced the capacity of MEFs to synthesize rRNA and to assemble ribosomes, explaining their reduced proliferation rate.

**Figure 5.**
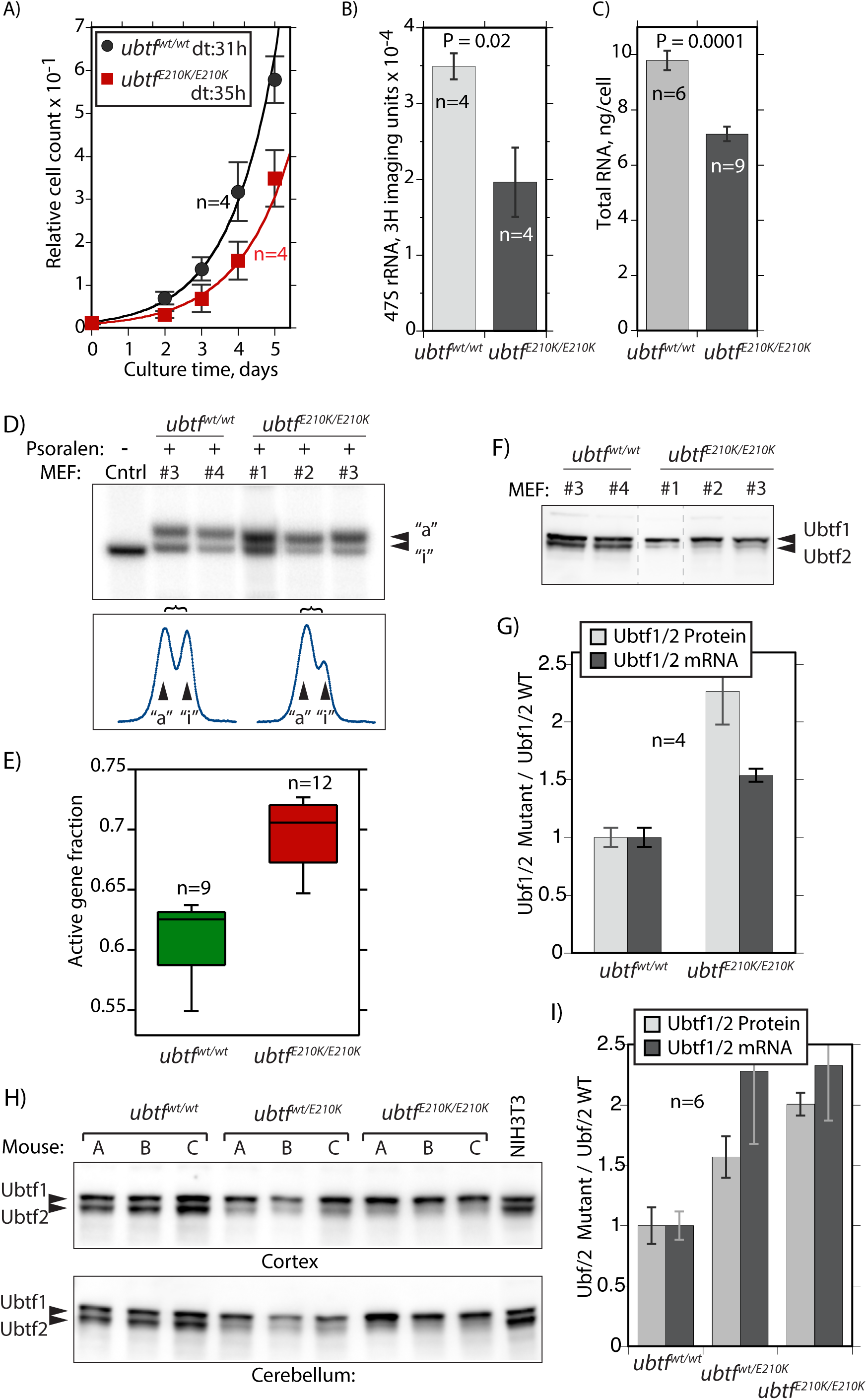
The E210K Ubtf mutation suppresses both proliferation and 47S pre-rRNA synthesis but enhances rDNA activation. A) *ubtf^E210K/E2109K^* MEFs were found to proliferate significantly more slowly than isogenic wild type (*Ubtf^wt/wt^*) MEFs. Doubling time (dt) was estimated from an exponential curve fit as respectively 35h and 31h for mutant and wild type MEFs. B) 47S pre-rRNA synthesis in *Ubtf^E210K/E2109K^* and wild type MEFs determined by metabolic pulse (30 min) labelling. See also Figure S9A for analysis of processing intermediates at increasing labelling times for individual MEF isolates. C) Per cell total cellular RNA content of *Ubtf^E210K/E2109K^* and wildtype MEFs. The data in A, B and C derive from two or more biological replicas in each of which a minimum of two independently isolated mutant and wild type MEF cultures were analyzed in parallel. D) PAC analysis of *Ubtf^E210K/E2109K^* and wild type MEFs. Upper panel shows an example of the active rDNA “a” and inactive “i” profiles for the 1.3kbp BamHI-BamHI 47S coding region fragment (see Figure 2A) and the lower panel corresponding band intensities. profiles. E) Active rDNA fractions were estimated from the combined curve fit analysis of 1.3, 2.4 and 4.7kbp BamHI-BamHI rDNA fragment PAC profiles. The data derive from three independent *Ubtf^E210K/E2109K^* and two wild type MEF isolates in two PAC biological replicas and are plotted to show median, upper and lower data quartiles and outliers. F) and G) Analysis of Ubtf1 and 2 levels in *Ubtf^E210K/E2109K^* and wild type MEF isolates. Panel F shows a typical Western analysis of Ubtf variants in these MEFs and panel G quantitative estimates of relative Ubtf1/Ubtf2 protein and mRNA ratios in these MEFs. H) and J) Show similar estimates of relative Ubtf1/Ubtf2 protein and mRNA ratios in Cortex and Cerebellum tissue from matched *Ubtf^wt/wt^*, *Ubtf^wt/E2109K^* and *Ubtf^E210K/E2109K^* adult mice. Error bars throughout indicate SDM.

### The E210K mutation also enhances Ubtf1 levels and the fraction of active rDNA repeats

Unexpectedly, the *Ubtf^E210K/E210K^* MEFs displayed a significant increase in the fraction of activated rDNA copies determined by PAC (Figure 5D and E), and this corresponded to an equally significant increase in the expression of the Ubtf1 variant both at the protein and mRNA levels (Figure 5F and G). A similar bias towards Ubtf1 expression was also observed in brain tissue of mutant mice (Figure 5H and I). This suggested the interesting possibility that the enhanced levels of Ubtf1 in the mutant MEFs revealed an inherent feedback mechanism regulating splicing. In this way the cell might control the fraction of active rDNA copies and hence potentially also rRNA synthesis. However, it will first be necessary to determine whether or not the E210K mutation directly affected usage of the adjacent splice junctions (see Figure S10B). In either scenario, the increase in active rDNA copies would normally be expected to enhance rRNA synthesis and cell growth in the mutant MEFs. Since this was clearly not the case, the E210K mutant MEFs displaying reduced rRNA synthesis, accumulation and proliferation (Figure 5A to C), we sought other origins for these effects.

### The E210K mutation reduces RPI loading and SL1 and Ubtf recruitment to the rDNA promoters

ChIP-qPCR analyses revealed that RPI loading across the rDNA was reduce by >40% in the *Ubtf^E210K/E210K^* mutant MEFs, explaining the observed reduction in pre-rRNA synthesis in these cells (compare RPI loadings in Figure 6A with *de novo* rRNA synthesis levels in 5B). Recruitment of Taf1B (SL1) and Ubtf to both Spacer and 47S rDNA promoters was somewhat reduced in the mutant MEFs, though less than RPI loadings (Figure 6B). Thus, the E210K Ubtf mutation most probably reduced pre-initiation complex formation, consistent with it affecting Ubtf-SL1 cooperation. The higher resolution of DChIP-Seq further showed that occupancy of Ubtf at both 47S and Spacer promoters was selectively reduced by the E210K mutation (Figure 6C), again consistent with a reduced Ubtf-SL1 cooperativity. The reduction of Ubtf at the rDNA promoters was particularly apparent in difference maps between wild type and E210K mutant MEFs (Figure 6D and S11). The reduction in Ubtf was especially strong at the Spacer promoter and corresponded with a similar reduction in Taf1B occupancy and in RPI recruitment ((Figure 6D and E). The data strongly suggested that the E210K mutation causes a small but significantly reduced ability of Ubtf to cooperate with SL1 in the formation of the RPI preinitiation complex, and together point to a reduction in the efficiency of RPI transcription initiation as the fundamental cause of the UBTF-E210K neuroregression syndrome. Thus, these data indirectly support the central role of the SL1-Ubf1 cooperation in determining RPI preinitiation complex formation and efficient rDNA transcription *in vivo*.

**Figure 6.**
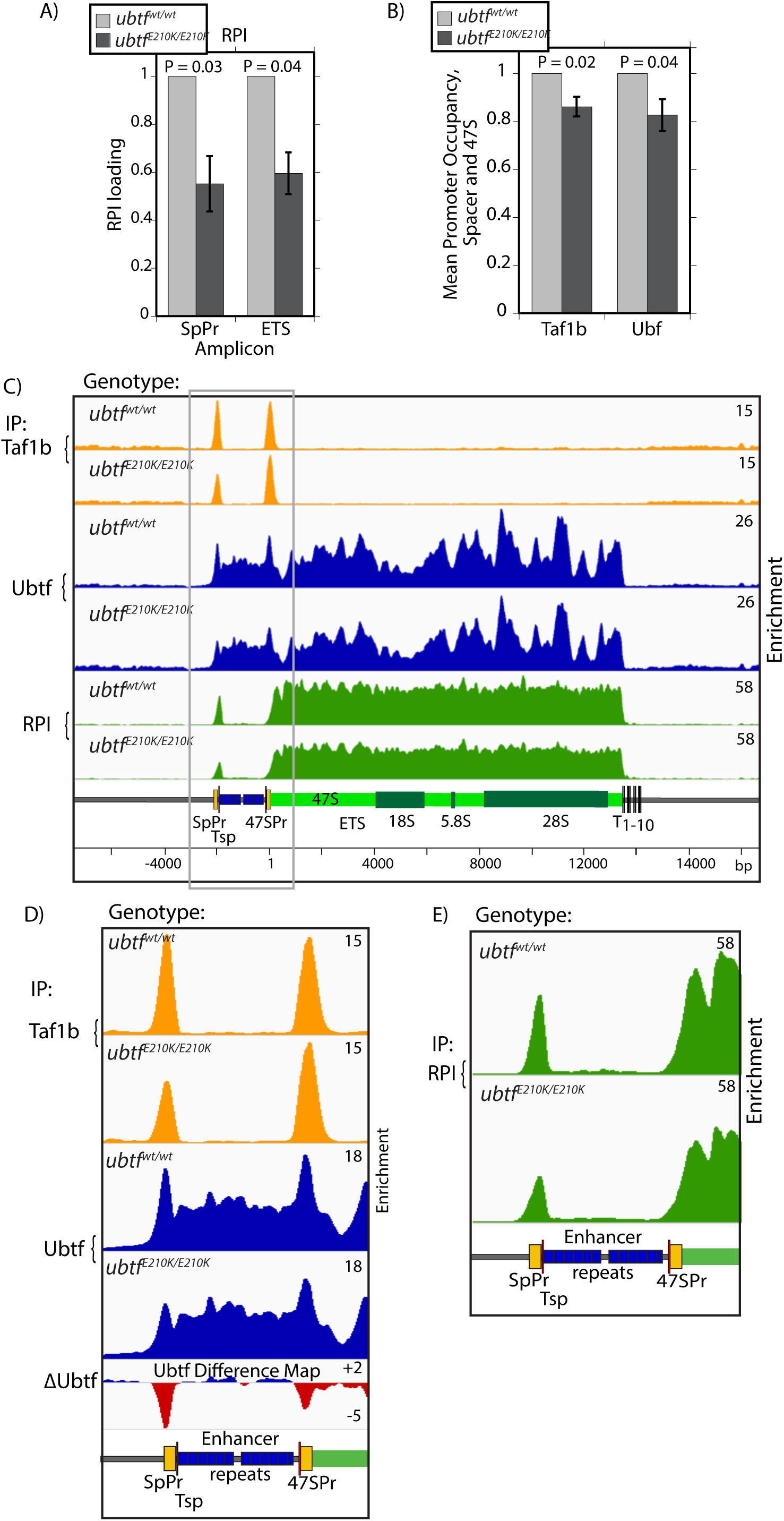
The E210K mutation in Ubtf suppresses both the RPI loading across the rDNA and preinitiation complex formation at the Spacer and 47S promoters. A) ChIP-qPCR analysis of RPI occupancy at the Spacer promoter within the ETS region of the 47S coding region in *Ubtf^E210K/E2109K^* and wild type MEFs. B) ChIP-qPCR analysis of relative preinitiation complex formation in *Ubtf^E210K/E2109K^* and wild type MEFs. The data show the mean occupancy at amplicons SpPr and T0/Pr by Taf1b or Ubtf. The data in A and B derive from 4 independent ChIP-qPCR experiments and error bars indicate the SEM, see Figure 2A and Materials and Methods for amplicon positions. C), D) and E) DChIP-Seq mapping of Taf1b, Ubtf and RPI across the rDNA of *Ubtf^E210K/E2109K^* and wild type MEFs. Panels D and E show an enlargement of the rDNA promoter and Enhancer region and a difference map of Ubtf occupancy (mutant - wild type MEFs). A full-width Ubtf difference map is shown in Figure S10. The data are typical of two biological replicas.

## DISCUSSION

Formation of the RPI preinitiation complexes on the rRNA genes (rDNA) in mammals determines as much as 35% of total nuclear RNA synthesis but is still poorly understood. In particular, prior to our study it was unclear how, or indeed if, the multi-HMGbox factor UBTF played a role in targeting the TBP-TAF_I_ complex SL1 to the rDNA promoters or if it simply acted as a general chromatin replacement protein. Ubtf displays little or no DNA sequence binding specificity and binds throughout the 15kbp NFR of the mouse and human rDNA (25). We previously showed that conditional deletion of the Ubtf gene inactivated rDNA transcription and allowed the reformation of nucleosomes across the rDNA (24, 26). Hence, it appeared that Ubtf may simply facilitate SL1 recruitment by eliminating the obstacles presented by nucleosomes. Genetic ablation of the SL1 subunit Taf1B has now revealed an unexpected sequence specific role for Ubtf and has suggested a novel induced-fit model for RPI preinitiation complex formation.

Though it was assumed from cell-free studies that SL1 would be essential for cell and organism survival due to its role in rDNA transcription, this had not been directly tested. Our data show that homozygous deletion of the gene for the SL1 subunit Taf1b prevented mouse development beyond blastocyst while heterozygous deletants were viable and fertile. Hence, as observed for the other RPI basal factors Ubtf and Rrn3, Taf1b is an essential factor in mouse. Conditional deletion of *taf1b* in MEF and mES cell culture was also found to arrest rDNA transcription and to cause severe disruption of nucleolar structure characteristic of nucleolar stress (24, 28). Depletion of Taf1b also prevented promoter recruitment of Taf1c and TBP subunits of SL1 and hence PIC formation at both the 47S pre-rRNA and the Enhancer-associated Spacer rDNA promoters. Quite unexpectedly, this also led to a loss of Ubtf at both these promoters, though not elsewhere across the rDNA NFR. ChIP-qPCR and high resolution DChIP-Seq showed that the loss of Ubtf from the promoters was proportional to the loss of SL1, strongly arguing that binding of these two basal factors was cooperative. Conversely, we had previously shown that *in cell* loss of Ubtf eliminated SL1 from the rDNA promoters (24, 26), consistent with the cooperative recruitment of these factors. Data from early cell-free studies had suggested two possible scenarios for RPI preinitiation complex formation, either SL1 recruitment depended on pre-binding of Ubtf or conversely that Ubtf recruitment depended on pre-binding of SL1 (22, 30). Our data resolve this contradiction by showing that in cells Ubtf and SL1 binding at the rDNA promoters is strongly interdependent, neither factor being recruited in the absence of the other. The lack of Ubtf binding at the promoters in the absence of SL1 was particularly surprising, especially so since Ubtf remained bound throughout the rest of the rDNA NFR and even at immediately promoter adjacent sites. Thus, the absence of SL1 the RPI promoters rather than being prefer sites of Ubtf binding as usually assumed, are quite on the contrary sites of low Ubtf affinity lying within the NFR continuum of higher affinity sites.

Our data further revealed the key importance of the Ubtf1 variant in the recruitment of SL1 to the rDNA promoters. Mouse and human cells express varying levels of the Ubtf1 and Ubtf2 splice variants that differ by a 37a.a deletion in HMGbox2 of Ubtf2 (Figure 4). By mapping these variants across the rDNA we found that Ubtf1 was recruited to the rDNA promoters at least four times more often than Ubtf2, though the data were also consistent with the exclusive recruitment of UBTF1 at the promoters. In contrast, Ubtf1 and Ubtf2 bound indistinguishably elsewhere across the rDNA. Since only Ubtf1 is present in mESCs, deletion of *taf1b* in these cells also clearly demonstrated that promoter recruitment of Ubtf1 depended on SL1 (Figures S4 and S5). Thus, formation of the RPI preinitiation complex is driven predominantly if not exclusively by a cooperation between SL1 and Ubtf1. This provides the first mechanistic explanation for why Ubtf1 is absolutely required for rDNA activity *in vivo* (31).

Recruitment of Ubtf1 and SL1 was found to be cooperative not only at the major 47S rDNA promoter but also at the enhancer associated Spacer promoter. Since these promoters display little DNA sequence homology, this raised the question of what in fact defines an RPI promoter and how is it recognized? Our data clearly show that promoter recognition involves the cooperative recruitment of Ubtf1 and SL1. Previous data showed that Ubtf interacts with SL1 solely via its highly acidic C-terminal tail, an ∼80 a.a. domain containing 65% Asp/Glu residues (32). However, this domain is not essential for cell-free transcription (33) and is anyhow present in both Ubtf variants. So, while it might play some role in bringing SL1 to the promoters it cannot explain their selective binding of Ubtf1. Co-immunoprecipitation also failed to detect any specific interaction between SL1 and one or other of the Ubtf variants (data not shown). Thus, it seems unlikely that the rDNA promoters are recognized by a pre-formed SL1-Ubtf1 pre-initiation complex. Rather we suggest that promoter recognition involves the transient imposition of a specific DNA conformation by Ubtf1 that is in turn locked into place by SL1 (Figure 7). There is significant precedent for such a mechanism, since the HMGboxes of Ubtf were shown to induce in-phase bending and looping of a DNA substrate. Indeed, it was suggested that such a looping could position UCE and Core promoter elements (Figure 1A) to facilitate their contact by SL1 (23, 34, 35). Essentially, Ubtf1 might transiently mould the promoter DNA to create an induced-fit for SL1, which it could then lock in place. Since the 37 a.a. deletion in HMGbox2 of Ubtf2 prevents this box from bending DNA (35), Ubtf2 would mould the promoter DNA differently from Ubtf1 and would therefore not induce the appropriate fit for SL1.

**Figure 7.**
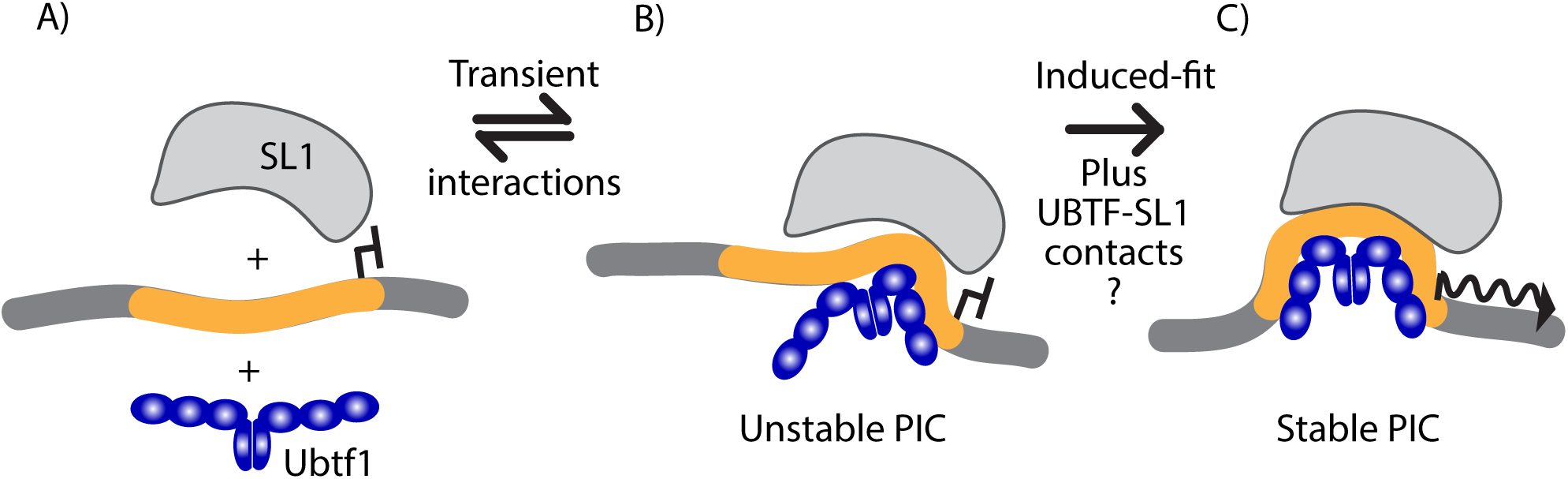
Diagrammatic representation of the induce-fit model for cooperative Ubft1-SL1 recognition and binding at the rDNA promoters. A) Neither Ubtf1 nor SL1 alone is able to form a stable interaction with the promoters. B) Ubtf1 binding induces a transient reshaping of the promoter that allows SL1 to form weak DNA interactions. C) Tightening of Ubtf contacts induces further promoter reshaping inducing a new DNA surface that closely “fits” the DNA interaction surface of SL1. The promoter and flanking DNA sequences are shown respectively in orange and dark grey, the region of Ubtf known to bend DNA is shown in blue, SL1 in light grey, and the active transcription initiation site is by an arrow.

Though in the induced-fit model of rDNA promoter recognition direct Ubtf1-SL1 contacts would not be essential, they may nonetheless play an important part in stabilizing the preinitiation complex. Indeed, our study of the UBTF-E210K recurrent pediatric neuroregression syndrome suggested that this was quite probably the case. Molecular modelling showed that while this E210K mutation in HMGbox2 of UBTF was very unlikely to affect interactions with DNA, it might well affect interactions with other proteins such as SL1. We found that introduction of the homozygous Ubtf-E210K mutation in MEFs significantly reduced rDNA transcription rates, reduce total cellular RNA accumulation and slowed cell proliferation. In apparent contradiction to these effects, the E210K mutation enhanced expression of Ubtf1 both in mutant MEFs and mouse tissues, and this led to an increase in the fraction of active rDNA copies, possibly as an attempt to compensate for reduced rDNA transcription. However, ChIP analyses further revealed that the Ubtf-E210K mutation reduced the cooperative recruitment of SL1 and Ubtf to the rDNA promoters. Thus, it appeared that the primary effect of the E210K-Ubtf mutation was to limit PIC formation on the rDNA. This further emphasized the central importance of a functional cooperation between Ubtf and SL1 in determining rDNA activity. It further suggested that the UBTF-E210K neurodegeneration syndrome was caused by a subtle defect in PIC formation on the rDNA.

In summary, our study identifies the parameters that determine RNA polymerase I promoter recognition and preinitiation complex formation *in vivo*. We reveal the central importance of a cooperative interaction between the RPI-specific TBP complex SL1 and the Ubtf1 splice variant in promoter recognition and propose an induce-fit model for pre-initiation complex formation that explains the functional differences between the Ubtf splice variants in terms of their abilities to induce specific conformational changes in the rDNA promoter sequences. Our data further suggest that the UBTF-E210K recurrent neurodegeneration syndrome is caused by a subtle reduction in UBTF-SL1 cooperativity that leads to reduced rDNA transcription.

## MATERIALS AND METHODS

### Primary antibodies for Immunofluorescence, ChIP and Western blotting

Rabbit polyclonal antibodies against mouse Ubtf, RPI large subunit (RPA194/Polr1A), and Taf1b were generated in the laboratory and have been previously described (24), anti-Taf1c was a gift from I. Grummt. All other antibodies were obtained commercially; anti-Fibrillarin (#LS-C155047, LSBio), anti-RPA135 (#SC-293272, Santa Cruz), anti-RPA194 (#SC-48385, Santa Cruz), anti-TBP (#ab818 and #ab51841, Abcam) and anti-FLAG M2 (F3165, Sigma).

### Generation of *taf1b* mutant mice

*Taf1b*+/- (targeted allele Taf1btm1a(EUCOMM)Hmgu) embryonic stem (ES) cells were obtained from EuMMCR and generated using the targeting vector PG00150_Z_6_E02. Two ES clones (HEPD0596_3_G02 and HEPD0596_3_H01) were each used to generate independent mouse lines using the services of the McGill Integrated Core for Animal Modeling (MICAM).

The resulting *taf1b^fl-neo^* mouse lines carried a “knock-out first” allele in which Lox recombination sites were inserted in intron 3 and intron 5, and a *neo* selective marker gene flanked by FRT sites inserted intron 3 (Figure S1A & B). Mouse lines heterozygous for the *taf1b^fl-neo^* allele derived from the two ES clones were viable, fertile, and appeared phenotypically normal compared to their wild-type littermate but no homozygotes were identified (data not shown). These mice were then crossed with FLPo (FLP recombinase) and Cre expressing mice (Jackson Laboratory strains FLPo (#012930), Sox2-Cre (#004783)) to generate both *taf1b^fl^* and *taf1b*^Δ^ alleles (Figure S1A). Subsequently Cre and FLPo transgenes were removed by backcrossing. *Taf1b^fl/fl^/ER-cre^+/+^*, *taf1b^fl/fl^/ER-cre^+/+^*/*p53^-/-^*and corresponding *taf1b^+/+^* control mouse lines were generated by crossing with ER-Cre expressing and p53-null mice (Jackson Laboratory strains ER-Cre (#004847) and p53 KO (#002101) and used to generate MEF and mES cell lines as previously described (24). Mice were genotyped by PCR using the primers (Figure S1A): A; 5’-gtcccttcctcactgatcac, B; 5’-tgcagattaggtggcctcag, C; 5’-ccctctcaccttctacccca and D; 5’-ctgggcttggtggctgtaa.

### Embryo collection and genotyping

Heterozygous *taf1b*^Δ/*wt*^ mice were inter-crossed and embryos isolated, imaged and genotyped from pregnant females at E3.5, 6.5, 7.5, 8.5 and E9.5 as described in (24, 26). DNA from E3.5 embryos was amplified using the REPLI-g Mini kit (QIAGEN). Individual embryos were genotyped by PCR using the same primers as for mouse lines (Figure S1A).

### Ethics statement concerning animal research

All animal care and animal experiments were conducted in accordance with the guidelines provided by the Canadian Council for Animal Protection, under the surveillance and authority of the institutional animal protection committees of Université Laval and the Centre hospitalier universitaire de Québec (CHU de Québec). The specific studies described were performed under protocol #2014-100 and 2014-101 examined and accepted by the “Comité de protection des animaux du CHU de Québec”. This ensured that all aspects of the work were carried out following strict guidelines to ensure careful, consistent and ethical handling of mice.

### Isolation and culturing of Taf1b conditional MEF and mES cells

Conditional Taf1b primary mouse embryonic fibroblasts (MEFs) were generated from E14.5 *taf1b^fl/fl^/ER-Cre^+/+^/p53^-/-^* and wild type control *ER-Cre^+/+^/p53^-/-^* embryos as previously described (26, 36), and were genotyped by PCR as described for mice, see Supplementary Data (Figure S1A and B).

Mouse Embryonic Stem (mES) cells were derived from the inner cell mass of *Taf1b^fl/fl^/ER-Cre^+/+^* and wild type control *ER-Cre^+/+^* blastocysts essentially as published (37). After establishment of the cells on feeder monolayers, they were adapted to feeder-independence on 2i/LIF N2B27 (ThermoFisher) free serum medium (38) and subsequently maintained in this medium. The *taf1b^fl/fl^/ER-Cre^+/+^* and control mESCs were genotyped as for MEFs.

Primary MEFs were also generated from E14.5 *ubtf^E210K/E210K^* and wild type control *ubtf^wt/wt^* sibling embryos from three independent litters, immortalized by transfection with pBSV0.3T/t (26) and genotyped by base sequencing of PCR products generated using primers 5’ CTGGGTGAAGTAGGCCTTGG and 5’ CCAGGAGGGTAAGGTGGAGA flanking the mutation site. All MEFs were cultured in Dulbecco’s modified Eagle medium (DMEM)-high glucose (Life Technologies), supplemented with 10% fetal bovine serum (Wisent, Life Technologies or other), L-glutamine (Life Technologies) and Antibiotic/Antimycotic (Life Technologies).

### Inactivation of *taf1b* in cell culture

Gene inactivation in MEF cultures followed the previously described procedures (24, 26). Briefly, cells were plated in 6 cm petri dishes (0.8×10^6^ cells each) and cultured for 18 hours in DMEM, high glucose, 10% fetal bovine serum or 2i/LIF N2B27 free serum medium as appropriate. For *taf1b* inactivation, 4-hydroxytamoxifen (4-HT) was added to both *taf1b^fl/fl^* and control cell cultures to a final concentration of 50nM (the 0h time point for analyses). After 4 hr incubation the medium was replaced with fresh medium without 4-HT. Cell cultures were then maintained for the indicated times and systematically genotyped by PCR on harvesting.

### Analysis of Taf1b and Ubtf1/2 protein and mRNA levels

Taf1b, and Ubtf1/2 protein levels were monitored by Western blotting. At harvesting, cells were quickly rinsed in cold phosphate buffered saline (PBS), recovered by centrifugation (2 min, 2000 r.p.m.) and resuspended directly in SDS–polyacrylamide gel electrophoresis (SDS-PAGE) loading buffer (39). After fractionation by SDS-PAGE, proteins were analysed by standard Western blotting procedures using an HRP conjugated secondary antibody and Immobilon chemiluminescence substrate (Millipore-Sigma). Membranes were imaged on an Amesham Imager 600 (Cytiva) and Ubtf1/2 ratios were determined from lane scans using ImageJ (40) and Gaussian curve fit using MagicPlot Pro (Magicplot Systems). Relative Ubtf1/2 mRNA levels were determined by PCR on total cDNA using primers bracketing the spliced sequences (5’TGCCAAGAAGTCGGACATCC and 5’TCCGCACAGTACAGGGAGTA). Products were fractionated by electrophoresis on a 1.5 or 2% agarose EtBr-stained gel, photographed using the G:BOX acquisition system (Syngene) and Ubtf1/2 mRNA ratios determined using ImageJ and Gaussian curve fitting as for proteins.

### Determination of rRNA synthesis rate

The rate of rRNA synthesis was determine by metabolic labelling immediately before cell harvesting. 10 µCi [^3^H]-uridine (PerkinElmer) was added per 1ml of medium and cell cultures incubated for a further 30min to 3h as indicated. RNA was recovered with 1 ml Trizol (Invitrogen) according to the manufacturer’s protocol and resuspended in Formamide (Invitrogen). One microgram of RNA was loaded onto a 1% formaldehyde/MOPS Buffer gel (41, 42) or a 1% formaldehyde/TT Buffer gel (43). The EtBr-stained gels were photographed using the G:BOX acquisition system (Syngene), irradiated in a UV cross-linker (Hoefer) for 5 min at maximum energy, and transferred to a Biodyne B membrane (Pall). The membrane was UV cross-linked at 70 J/cm^2^, washed in water, air dried and exposed to a Phosphor BAS-IP TR 2025 E Tritium Screen (Cytiva). The screen was then analyzed using a Typhoon imager (Cytiva) and quantified using the ImageQuant TL image analysis software.

### Psoralen crosslinking accessibility and Southern blotting

The psoralen crosslinking accessibility assay and Southern blotting were performed on cells grown in 60 mm petri dishes and DNA was analyzed as previously described (44, 45), using the 6.7kb 47SrRNA gene EcoRI fragment (pMr100) (44). The ratio of “active” to “inactive” genes was estimated by analyzing the intensity profile of low and high mobility bands revealed by phospho-imaging on an Amersham Typhoon (Cytiva) using a Gaussian peak fit generated with MagicPlotPro (MagicPlot Systems LLC).

### Indirect immunofluorescence (IF) microscopy

Cells were plated on poly-lysine treated coverslips and subjected to the standard 4-HT treatment to induce *taf1b* deletion. At the indicated time points cells were rinsed with PBS, fixed in 4% PFA, PBS for 10 min and permeabilized with 0.5% Triton, PBS for 15 minutes. After a blocking step in PBS-N (PBS, 0.1% IGEPAL (Sigma)), 5% donkey serum, cover slips were incubated with primary antibodies in PBS-N, 5% donkey serum for ∼16 h at 4 deg. C. RPI was detected using a combination of mouse anti-A194 and A135 antibodies (#SC-293272, #SC-48385), fibrillarin with goat anti-FBL (#LS-C155047), and Ubtf with rabbit anti-Ubtf (in-house #8). Cells were incubated for ∼ 2 h at room temperature with the appropriate AlexaFluor or Dylight 488/568/647 conjugated secondary antibodies (ThermoFisher / Jackson ImmunoResearch) and counterstained with DAPI. After mounting in Prolong Diamond (ThermoFisher), epifluorescent 3D image stacks were acquired using a Leica SP5 II scanning confocal microscope and LAS-AF (Leica Microsystems) and Volocity (Quorum Technologies) software.

### Chromatin immunoprecipitation (ChIP)

Cells were fixed with 1% formaldehyde for 8 min at room temperature. Formaldehyde was quenched by addition of 125 mM Glycine and cells harvested and washed in PBS. Nuclei were isolated using an ultrasound-based nuclei extraction method (NEXSON: Nuclei Extraction by SONication) (46) with some modifications. Briefly, for all cell types, 33 million cells were resuspended in 1.5 ml of Farnham lab buffer (5 mM PIPES pH 8.0, 85 mM KCl, 0.5% IGEPAL, protease inhibitors). Cell suspensions were sonicated in 15 ml polystyrene tubes (BD #352095) using 3 to 4 cycles of 15 sec on : 30 sec off at low intensity in a Bioruptor (Diagenode). After recovery of the NEXSON-isolated nuclei by centrifugation (1000g, 5 min), nuclei were resuspended in 1.5 ml of shearing buffer (10 mM Tris-HCl pH 8.0, 1 mM NaEDTA, 0.1% SDS, protease inhibitors) and sonicated for 25 min, 30 sec on : 30 sec off, at high intensity. Each immunoprecipitation (IP) was carried out using the equivalent of 16 x 10^6^ cells as previously describe (24). To map Ubtf1 and Ubtf2 variants cDNAs encompassing the complete coding regions (M61726 / M61725) were subcloned into pCDNA3-3xFLAG-C1 (N. Bisson) and verified by Sanger sequencing. The resulting pC3xFLAG-UBF1 and -UBF2 (lab. stocks #2072, #2073) constructs were used to transfect NIH3T3 and cell cultures selected with G418. Pools of positive clones expressing Ubtf1 or Ubtf2 at sub-endogenous levels were then subjected to parallel ChIP for FLAG-Ubtf1/2 (anti-FLAG) and total Ubtf.

### ChIP-qPCR analysis

All ChIP experiments included a minimum of 2 biological replicates and were analyzed as previously described (24). For qPCR analysis, reactions (20 μl) were performed in triplicate using 2.5 μl of sample DNA, 20 pmol of each primer, and 10 μl of Quantifast SYBR Green PCR Master Mix (QIAGEN) or PowerUp™ SYBR™ Green Master Mix (ThermoFisher). Forty reaction cycles of 10 s at 95 and 30 s at 58 °C were carried out on a Multiplex 3005 Plus (Stratagene/Agilent). The amplicon coordinates relative to the 47S rRNA initiation site (BK000964v3) were as follows: IGS3, 42653-42909; SpPr, 43076-43279; Tsp, 43253-43447; T0/Pr (47SPr), 45133-40; 47S-5’, 159-320; ETS, 3078-3221; 28S, 10215-10411; T1-3, 13417-13607. Data was analyzed using the MxPro software (Agilent). The relative occupancy of each factor was determined by comparison with a standard curve of amplification efficiency for each amplicon using a range of input DNA amounts generated in parallel with each qPCR run.

### ChIP-Seq and data analysis

ChIP DNA samples were quality controlled by qPCR before being sent for library preparation and 50 base single-end sequencing on an Illumina HiSeq 2500 or 4000 (McGill University and Genome Quebec Innovation Centre). Sequence alignment and deconvolution of factor binding profiles to remove sequencing biases (Deconvolution ChIP-Seq, DChIP-Seq) were carried out as previously described (9, 24). The manual for the deconvolution protocol and a corresponding Python script can be found at https://github.com/mariFelix/deconvoNorm. Gaussian curve fitting to transcription factor binding profiles was perform using MagicPlot Pro (Magicplot Systems) on data from the DChIP-Seq BedGraph files. The raw sequence files and the processed deconvolution BedGraphs have been submitted to ArrayExpress under accession E-MTAB-10433.

### Cell Proliferation Assay

Cells from two *ubf^wt/wt^* MEF clones (#3, #4) and three *ubf^E210K/E210K^* MEF clones (#1, #2, #3) were continuously cultured for more than a week prior to assay. Cells were plated at ∼500 per well in 96-well plates and cultured for six days. At each timepoint, duplicate wells were treated with Hoechst 3342 (Invitrogen, Thermo Fisher Scientific) for 45min. Images were acquired using Cytation5 (Cell Imaging Multi-Mode Reader by BioTek) and cell counts for each clone were determined using the Gen5 software.

### Total RNA Extraction and Quantification

Cells were trypsinized, counted and total RNA was recovered from 3×10^6^ cells using 1 ml of Trizol (Invitrogen, Thermo Fisher Scientific) according to the manufacturer’s protocol. RNA yields were determined using Qubit RNA BR (Invitrogen, Thermo Fisher Scientific).

## FUNDING

This work was supported by operating grants from the Canadian Institutes of Health Research (CIHR), [grant number MOP12205/PJT153266] and the Natural Science and Engineering Council (NSERC) of Canada [grant number RGPIN-2017-06128] to T. Moss, and the National Institutes of Health of the U.S.A. (NIH) [grant numbers R21 GM118962 and R03 NS114616] respectively to R. T. Hori and M. M. Khan. The Research Centre of the Québec University Hospital Centre (CRCHU de Québec-Université Laval) is supported by the Fonds de Recherche du Québec - Santé (FRQS).

## ACKNOWLEDGEMENTS

We would like to thank the staff of the “Centre d’Expertise et de Services Génome Québec” for Next-Generation Sequencing, Marianne Sabourin-Felix (Cancer Division, CRCHU de Québec-Université Laval) for sequence alignment and DChIP-Seq deconvolution and Mark Robinson and Helen Lindsay (IMLS/SIB, University of Zürich) for making their computing facilities and advice available to us.

## SUPPORTING INFORMATION CAPTIONS

### SUPPORTING RESULTS

#### The Taf1B gene is essential for mouse development beyond early blastula

Mouse lines carrying a targeted “Knockout First” insertion in the gene for Taf1B (Taf68), were established and these crossed to remove the *ß-Gal* and *Neo* cassette insertion, generating lines carrying lox sites flanking exons 4 and 5 of *taf1b* (Figure S1A and B). Subsequent recombination of these lox sites inactivated the *taf1b* gene, (Figure S1C), see Supplementary Materials and Methods for more detail. Mice heterozygous for the *taf1b*^Δ^ allele were found to be both viable and fertile and the null-allele was propagated at near Mendelian frequency (Table S1). However, no *taf1b*^Δ/Δ^ homozygous offspring (pups) were identified and genotyping of embryos detected no *taf1b*^Δ/Δ^ homozygotes at stages 6.5 and later. In contrast, four *taf1b*^Δ/Δ^ embryos were detected at 3.5 dpc, though only one of these displayed a recognizable blastula morphology (Figure S1D and E). It was concluded that *taf1b* was essential for mouse development beyond blastula but that maternal Taf1B mRNA or protein, or simply ribosome availability may have been sufficient to support development beyond the morula stage. This is fully consistent with the previous data for inactivation of the TBP gene *tbp*/*gtf2d* (47), and suggests the interesting possibility that the effects of TBP-loss on early development could in large part be due to inactivation of RPI transcription. In support of this possibility, the SL1 complex is known to be generally less abundant than the RPII/PolII TFIID complex (48) and so could be limiting for embryo growth. Further, inactivation of the genes for the RPI factors Ubtf (*ubtf)* and Rrn3/TIF1A (*rrn3*) arrest mouse development during early cleavage stages (24, 26). A similar argument could be made for inactivation of RPIII/PolIII transcription since loss of the Brf1 subunit of the TFIIIB complex also causes developmental arrest during early cleavage stages (49). We conclude that the maternal protein translation machinery is limiting in the cleavage embryo and must be replenished by zygotic expression to allow further development.

### SUPPORTING TABLE

**Table S1.** Numbers and genotypes of embryos and pups derived from matings of *Taf1b+/-* mice.

| Age (dpc) | Total No. | *Taf1b*<sup>wt/wt</sup> | *Taf1b*<sup>Δ/wt</sup> | *Taf1b*<sup>Δ/Δ</sup> |
| --- | --- | --- | --- | --- |
| 3.5 | 20 | 3 (15%) | 13 (65%) | 4 (20%) |
| 6.5 | 23 | 12 (52%) | 11 (48%) | 0 (0%) |
| 7.5 | 15 | 6 (40%) | 9 (60%) | 0 (0%) |
| 8.5 | 35 | 8 (23%) | 27 (77%) | 0 (0%) |
| 9.5 | 16 | 4 (25%) | 12 (75%) | 0 (0%) |
| Pups | 112 | 35 (31%) | 77 (69%) | 0 (0%) |

### SUPPORTING FIGURE LEGENDS

**Figure S1.**
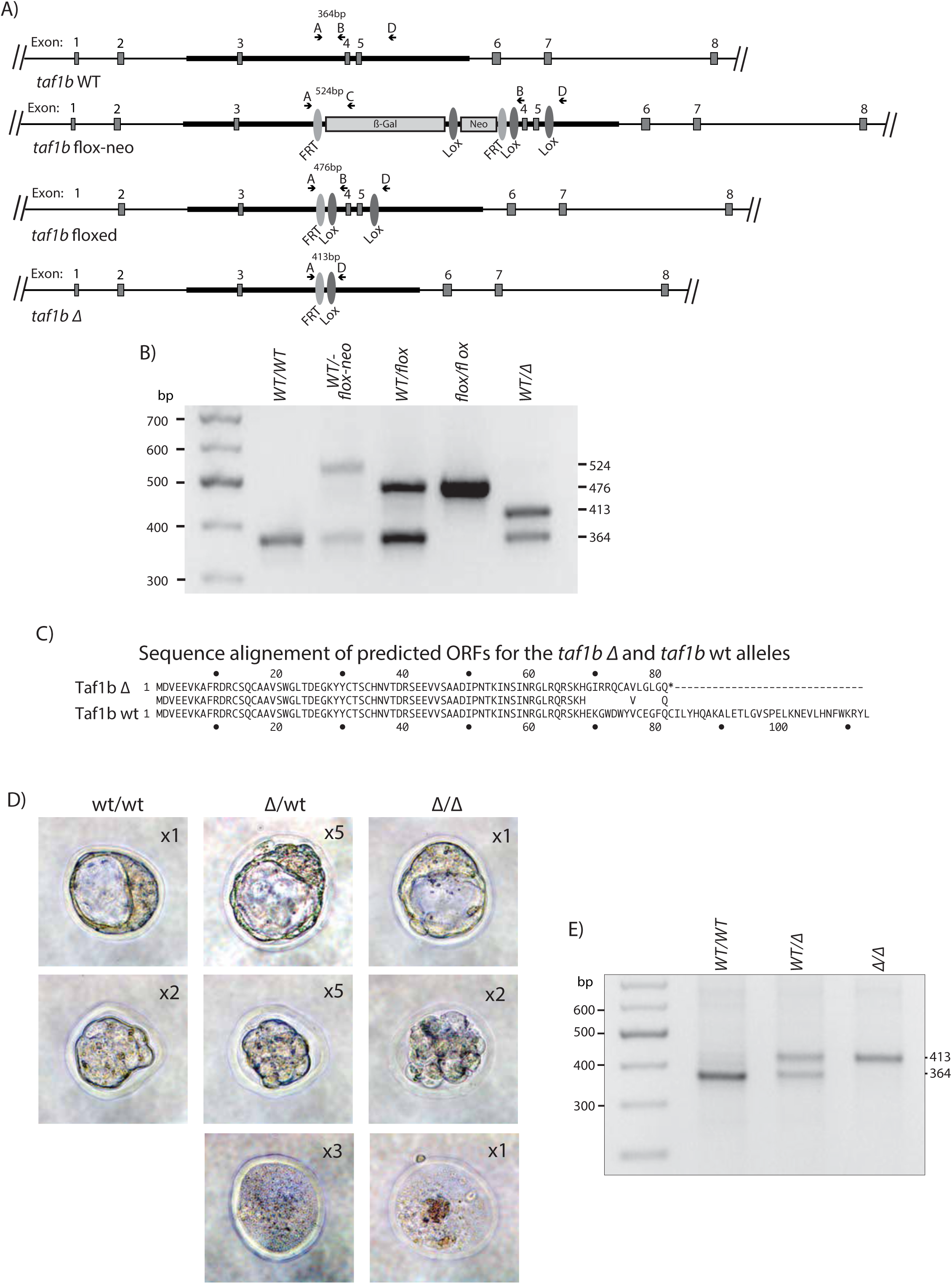
Construction and phenotypic effects of the mutant *taf1b* alleles A) Organisation of the first 8 exons of the mouse *taf1b* gene (*taf1bwt)*, and the “flox-neo” insertion, “floxed” and alleles indicating the position of inserted FRT and Lox sites and the inactivated (*taf1b*Δ) allele after Lox site recombination to delete exons 4 and 5. The positions of genotyping primers A to D are also indicated. B) Examples of mouse PCR genotyping. C) alignment of the N-terminal sequence of wild type Taf1b with the predicted residual Taf1b peptide encoded by the *taf1b*Δ allele. D) Typical images of mouse embryos at 3.5 dpc derived from *taf1b*^*wt*/Δ^ mouse crosses. The corresponding genotypes and numbers of embryos in each class are indicated, see also Table S1. E) Embryo phenotyping using primers A and D shown in panel A.

**Figure S2.**
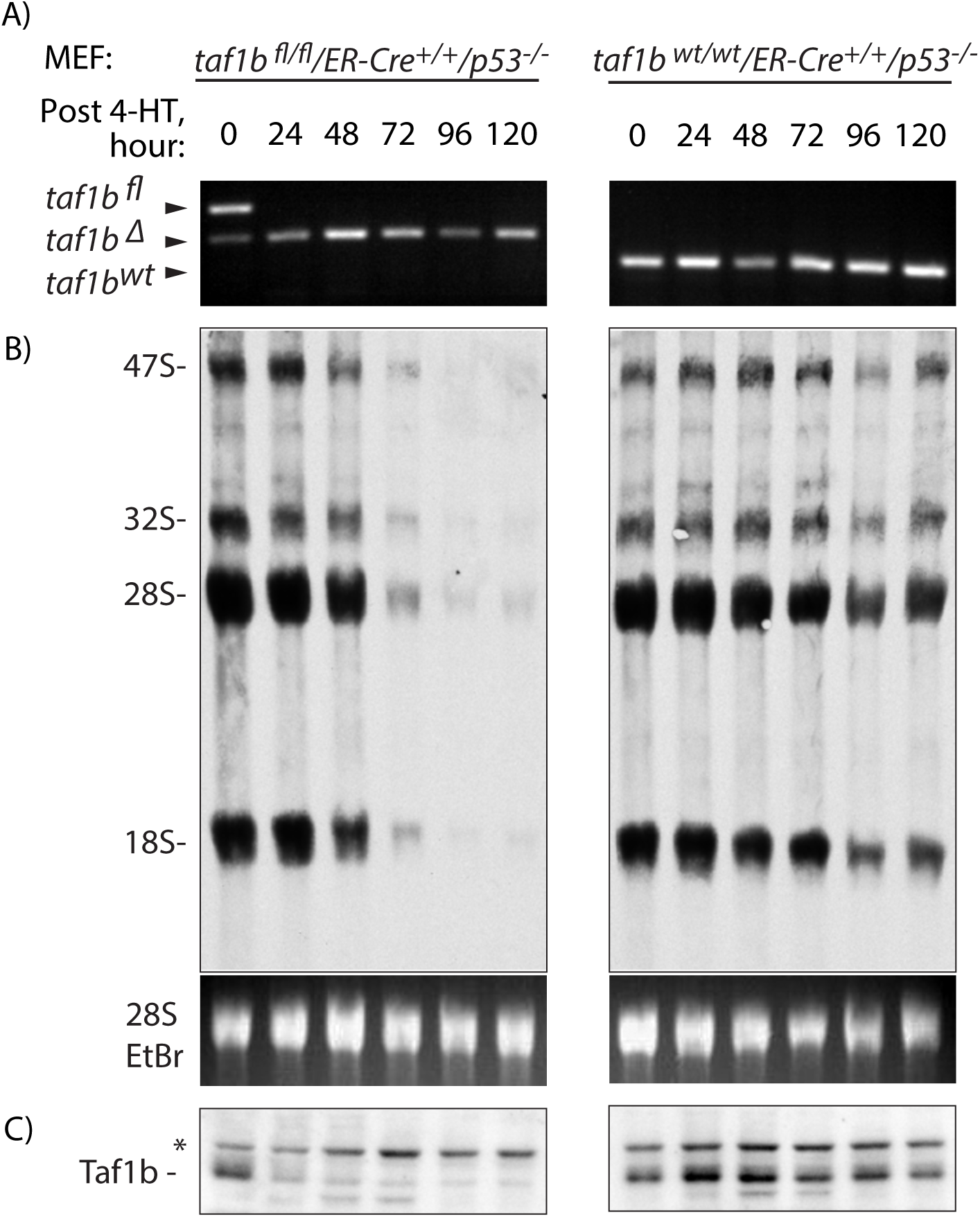
Time course of rRNA synthesis after *taf1b* inactivation in MEFs. Conditional *taf1b^fl/fl^/p53^-/-^/ERcre^+/+^* and control *taf1b^wt/wt^/p53^-/-^/ERcre^+/+^* MEFs were treated with 50 nM 4-hydroxy-tamoxifen (4-HT) for 4 h (4-HT pulse) before removing 4-HT by a change of the culture medium. 47S pre-rRNA synthesis was then determined by [3H]-uridine RNA metabolic labelling. The panels show the parallel time course analyses post 4-HT treatment of; A) *taf1b* inactivation, B) [3H]-rRNA labelling versus steady state 28S rRNA, and C) Taf1b protein depletion. The 47S pre-rRNA and rRNA processing products are indicated in B. The “*” in C indicates a non-specific antibody interaction that serves as a loading control.

**Figure S3.**
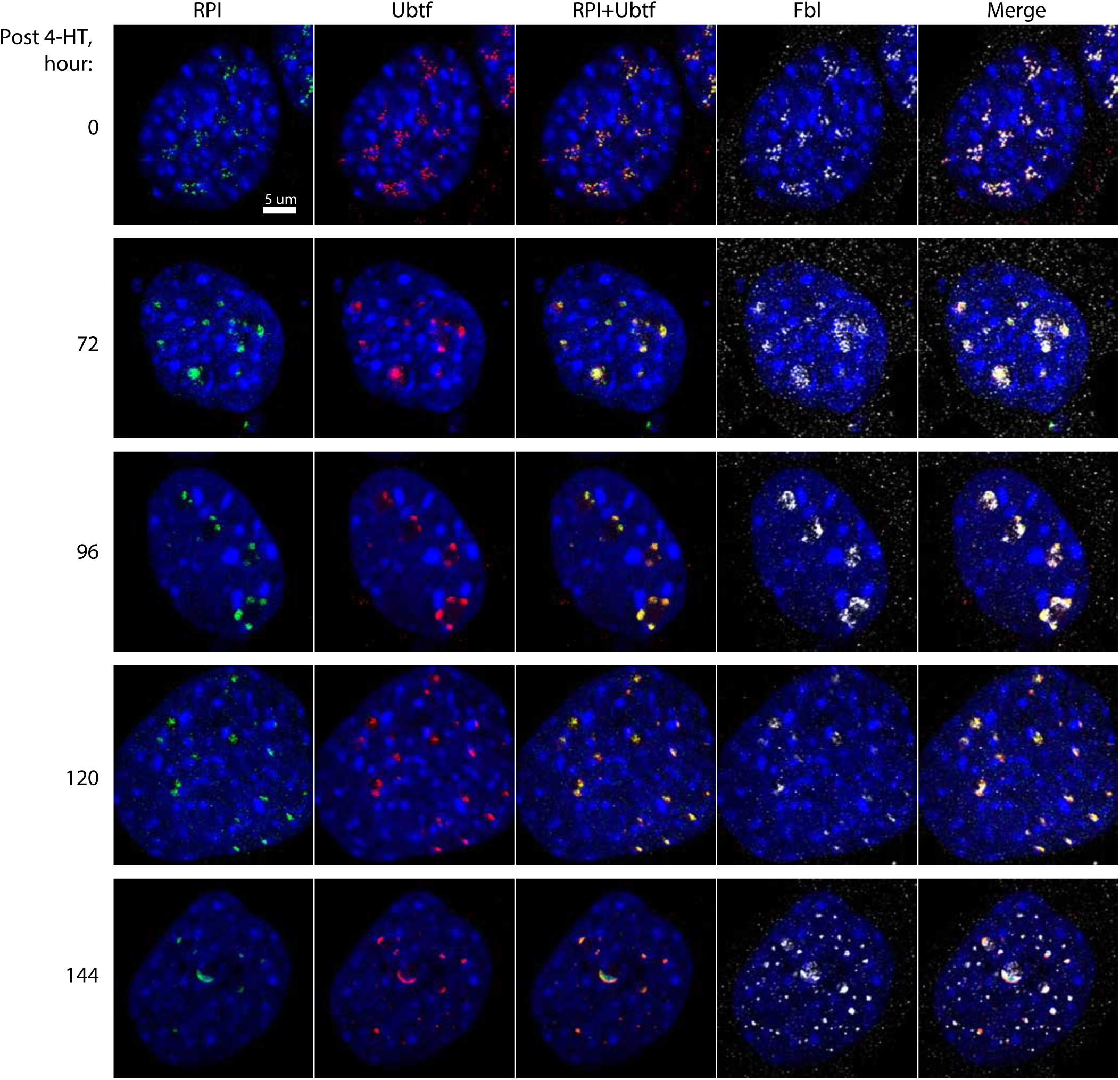
Time course of parallel *in situ* immunofluorescence labelling of RPI, Ubtf and fibrillarin (Fbl) in *taf1b^fl/fl^/p53^-/-^/ERcre^+/+^* MEFs after 4-HT induction of *taf1b* inactivation. Single confocal image planes are shown as for each factor and as merged overlays each with DAPI staining of DNA.

**Figure S4.**
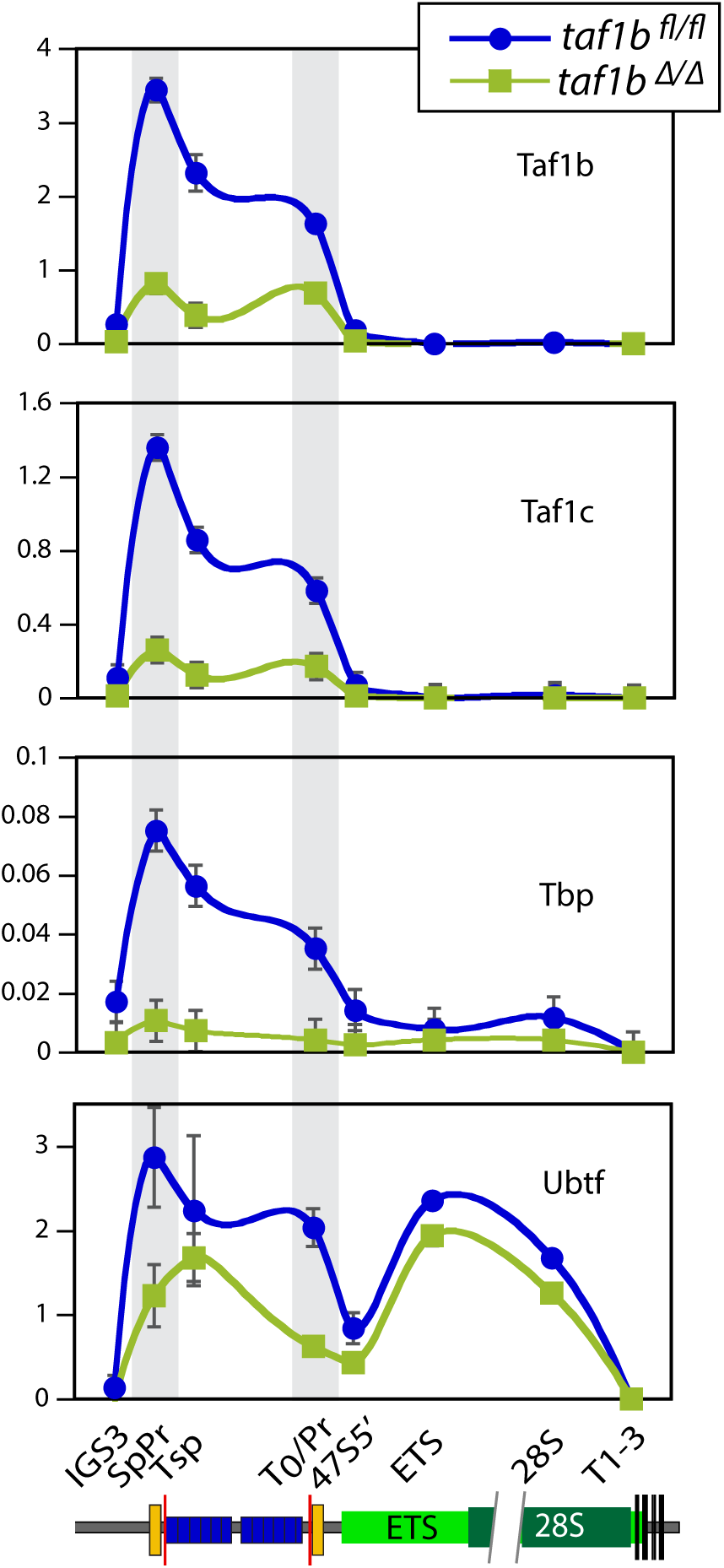
ChIP-qPCR analysis of Taf1b, Taf1c, Tbp and Ubtf occupancy at sites across the rDNA of *taf1b^fl/fl^/ERcre^+/+^* mESCs before and after *taf1b* inactivation by 4-HT treatment. The data for Taf1b and Ubtf derive from 3 ChIP biological replicas and for Taf1c and Tbp a single ChIP each analyzed by qPCR in triplicate.

**Figure S5.**
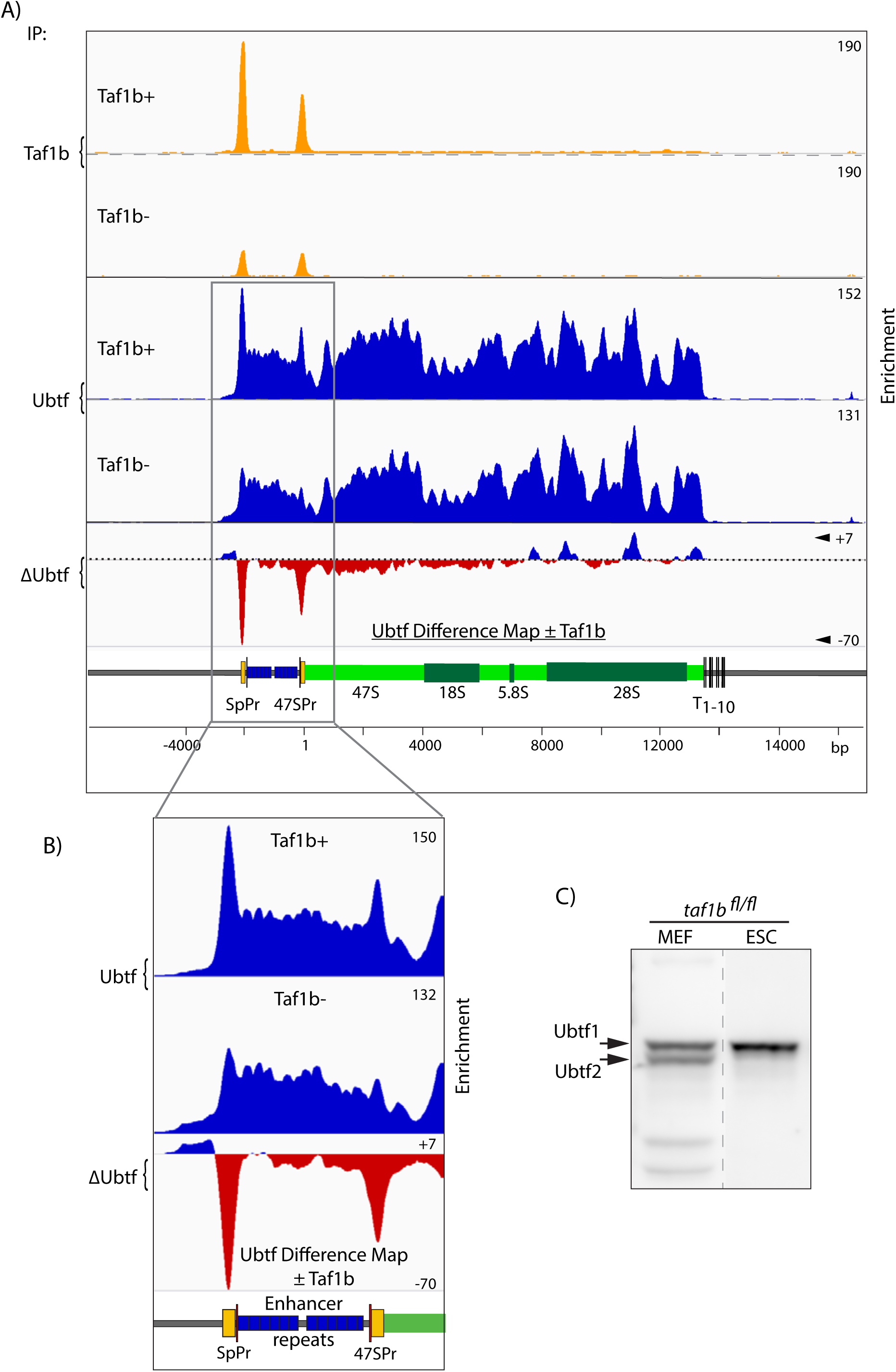
Taf1b depletion in condition *taf1b^fl/fl^/ERcre^+/+^* mESCs also induces depletion of Ubtf from both Spacer and 47S promoters but not from the adjacent enhancer repeats or from the 47S gene body. A) DChIP-Seq analysis of Taf1b and Ubtf occupancy across the rDNA repeat before (Taf1b+) and 3 days after *taf1b* inactivation (Taf1b-). ΔUbtf indicates the difference map of Ubtf occupancy after Taf1b depletion minus the occupancy before Taf1b depletion. B) Magnified view of the DChIP mapping in A showing detail over the promoter and enhancer regions. C) Comparative Western blots of Ubtf from Taf1b conditional MEFs (*taf1b^fl/fl^/p53^-/-^/ERcre*^+/+^) and mESCs (*taf1b^fl/fl^/ERcre^+/+^*), showing the presence of both Ubtf1 and Ubtf2 variants are expressed in the MEFs but only the Ubtf1 variant is expressed in the mESCs.

**Figure S6.**
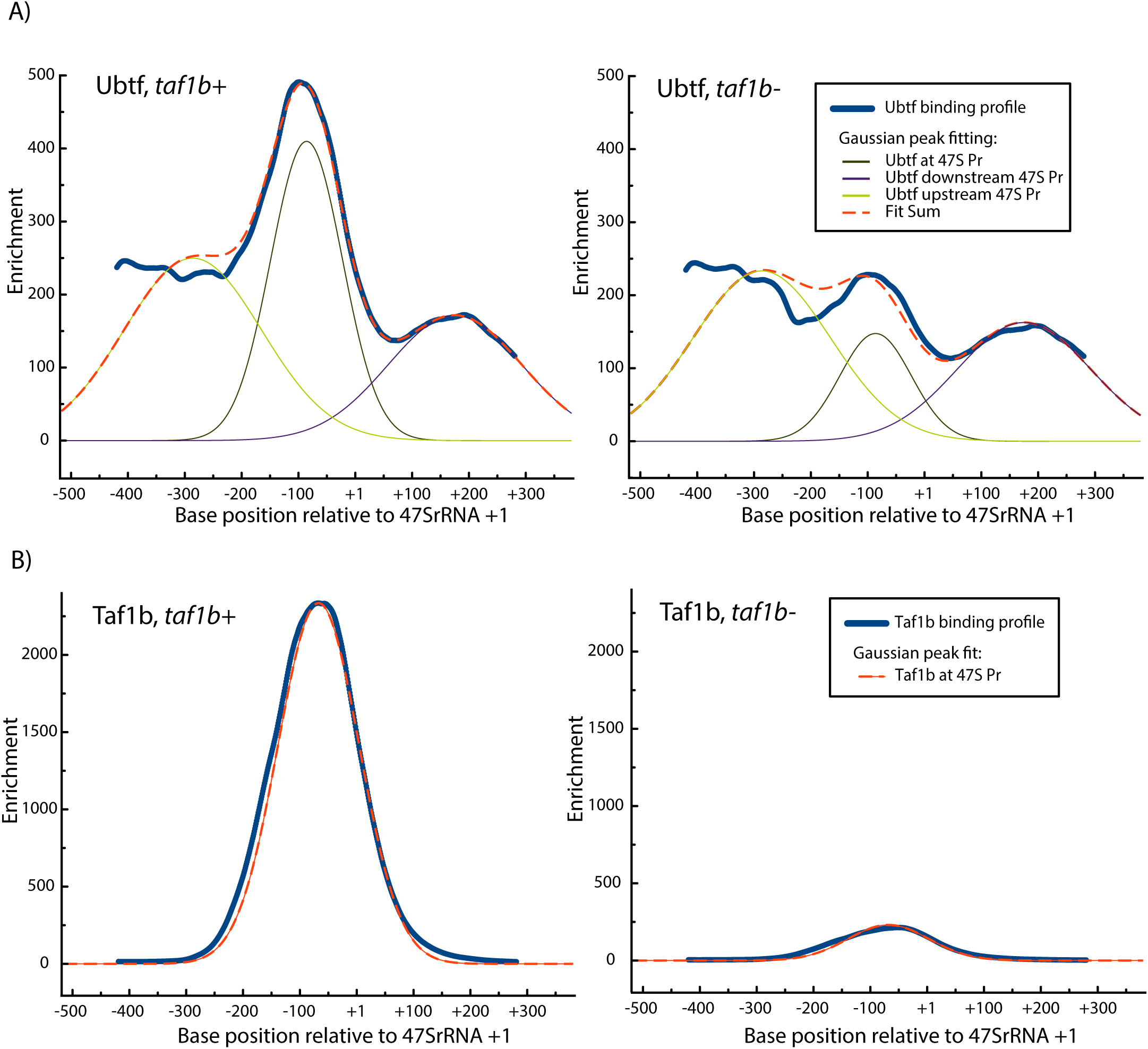
Analysis of DChIP profiles by Gaussian curve fit before and after *taf1b* inactivation. A) and B) respectively show examples of Ubtf and Taf1b enrichment profiles over the 47S promoter region before and after *taf1b* inactivation are shown, (dark blue line), and the best Gaussian peak fits to these profiles (dashed red line). In the case of Taf1b the profile closely followed a single Gaussian peak from which both the position and relative occupancy were determined. Since Ubtf was present not only at the promoter but also over the adjacent regions, curve fits were made using three Gaussians peaks, and the central one used to estimate relative occupancy.

**Figure S7.**
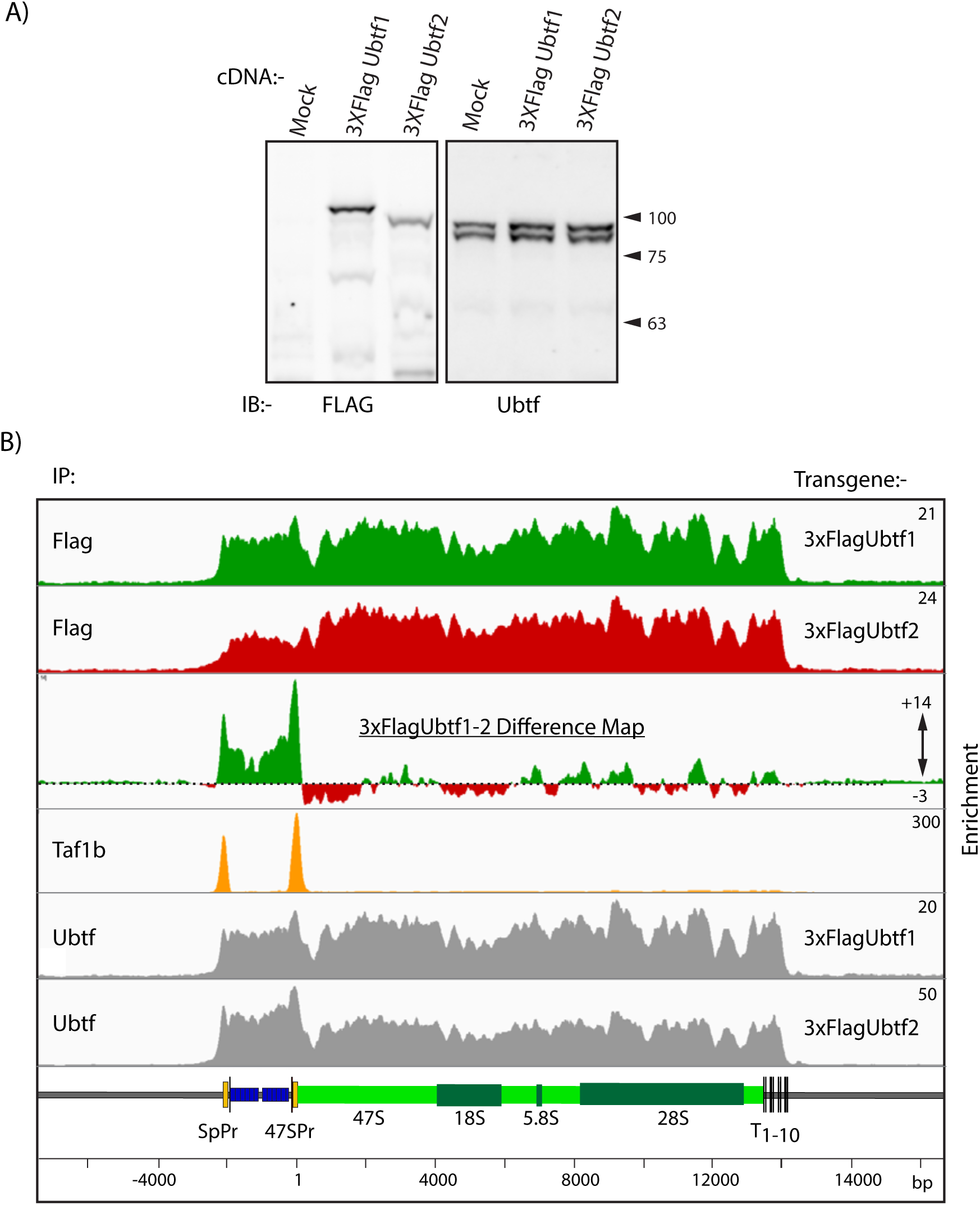
Mapping of Ubtf1 and −2 variants across the rDNA unit. A) Expression of exogenous 3xFlag-Ubtf1 and Ubtf2 in NIH3T3 MEFs. Total cell protein extracts from Mock, 3xFlag-Ubtf1 or 3xFlag-Ubtf2 transfected cells were analyzed by Western blot using either anti-Flag (αFlag), left hand panel) or anti-Ubtf (Ubtf) antibodies to detect total Ubtf (right hand panel). B) DChIP-Seq mapping profiles of the exogenously 3xFlag-Ubtf1 or 3xFlag-Ubtf2 (Flag) expressed in NIH3T3 MEFs (as in Figure 4) and the total endogenous Ubtf profiles ChIPped from the same chromatin preparations.

**Figure S8.**
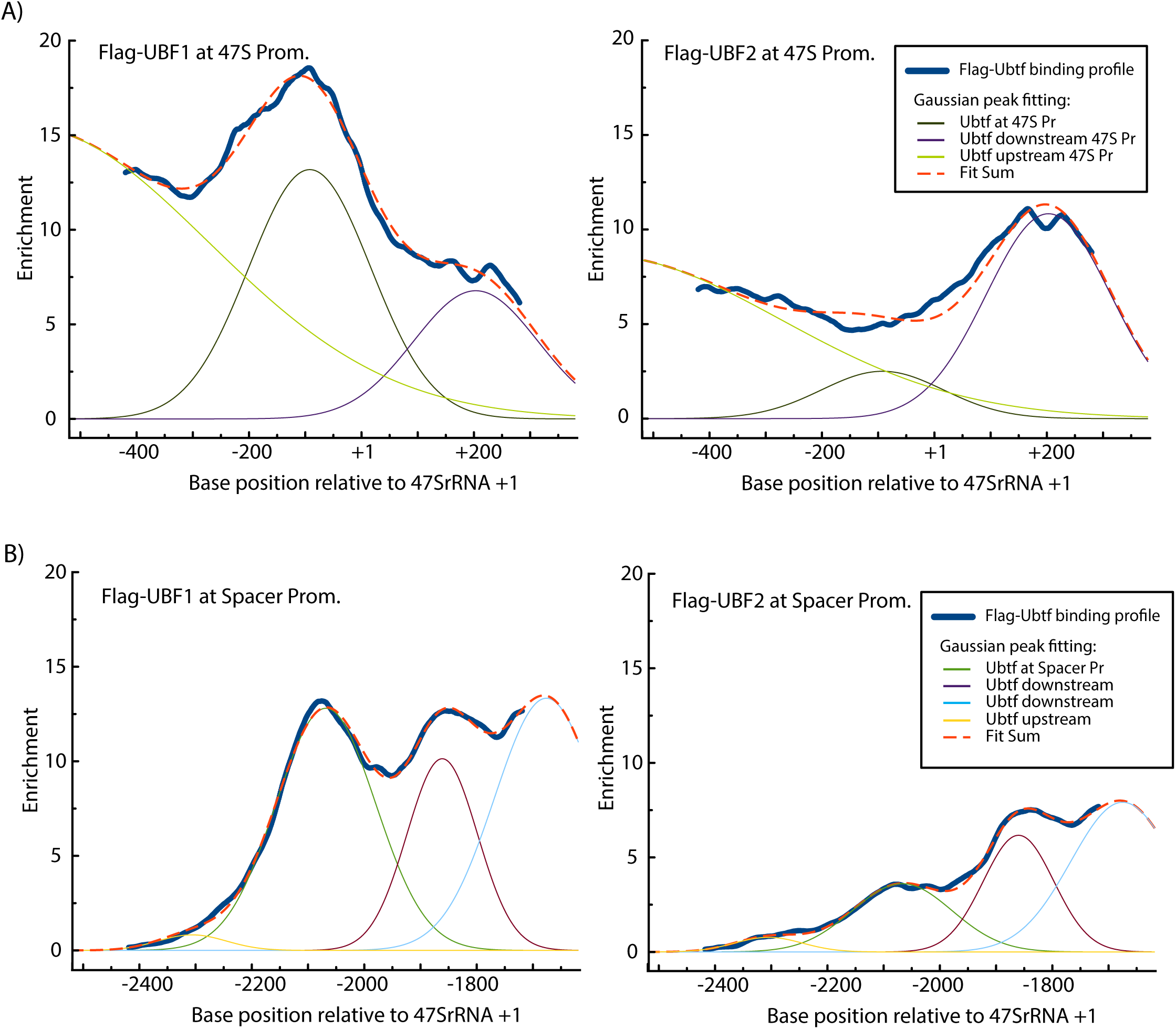
Analysis of DChIP profiles for Flag-Ubtf1 and -Ubtf2 by Gaussian curve fit. A) and B) Show examples of Flag-Ubtf1 and Flag-Ubtf2 enrichment profiles respectively over the 47S and Spacer promoter regions, (dark blue line). The best Gaussian peak fits to these profiles are shown (dashed red line), as are the individual Gaussian peaks used to estimate relative promoter occupancy. Since Ubtf was present not only at each promoter but also over adjacent regions, curve fits were made using three, or in the case of the Spacer promoter four, Gaussians peaks.

**Figure S9.**
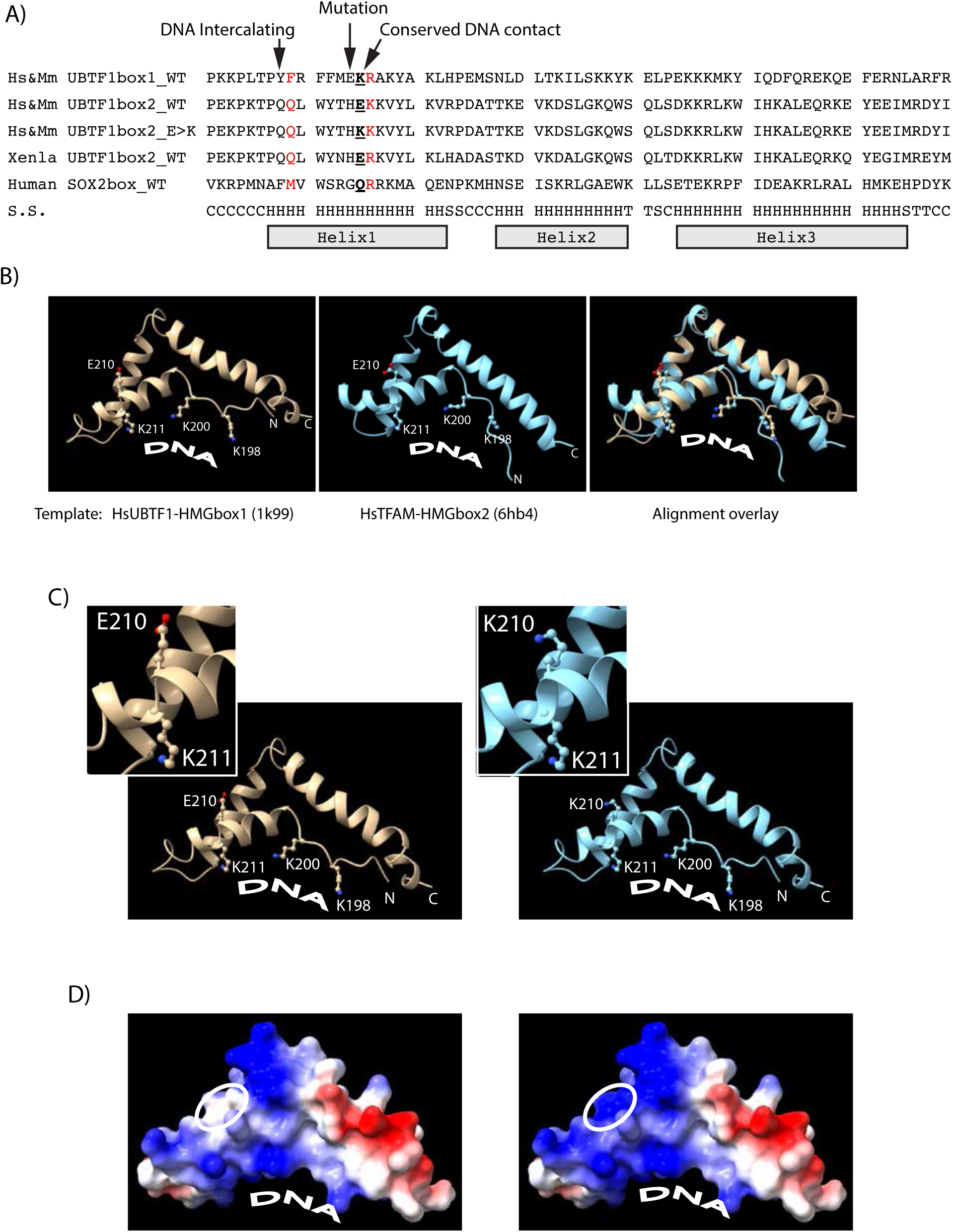
Structure prediction for wild type and E210K mutant HMGbox2 of Ubtf1. A) Sequence alignment of the HMGboxes 1 and 2 of human and mouse Ubtf1 (NM_014233-2, NP_035681) with HMGbox2 of Xenopus laevis Ubtf1a (CAA42523.1) and the HMGbox of SOX2 (P48431). The positions of the predicted DNA intercalating residue, the E210K mutation and the adjacent conserved basic DNA contacting residue are indicated as are the positions of the α-helical segments. B) Comparative molecular modelling of Ubtf HMGbox2 using as templates the structures 1k99 (human Ubtf HMGbox1) and 6hb4 (human mitochondrial transcription factor A, TFAM). The two predicated structures were generated by SWISS-MODEL (50) and are shown individually and as an aligned overlay generated in ChimeraX-1.1.1 (51). Comparison of these structures using the Matchmaker routine in ChimeraX-1.1.1 revealed an RMSD of 1.215 Å over 41 of 72 alpha-carbons, including those of helix 1 affected by the E210K mutation. C) The predicted positions and orientations of the E210 and K210 residues within the HMGbox2 of Ubtf1 are shown relative to the adjacent conserved basic residue at position 211, which is a lysine in UBTF/Ubtf. The likely other DNA minor groove contacting residues K198 and K200 are also shown. D) The predicted surface electrostatic potential of the wild type, left, and the mutant, right, HMGbox2. Blue indicates a positive and red a negative potential. Position of changes in surface potential due to the E210K mutation are enclosed by an ellipse.

**Figure S10.**
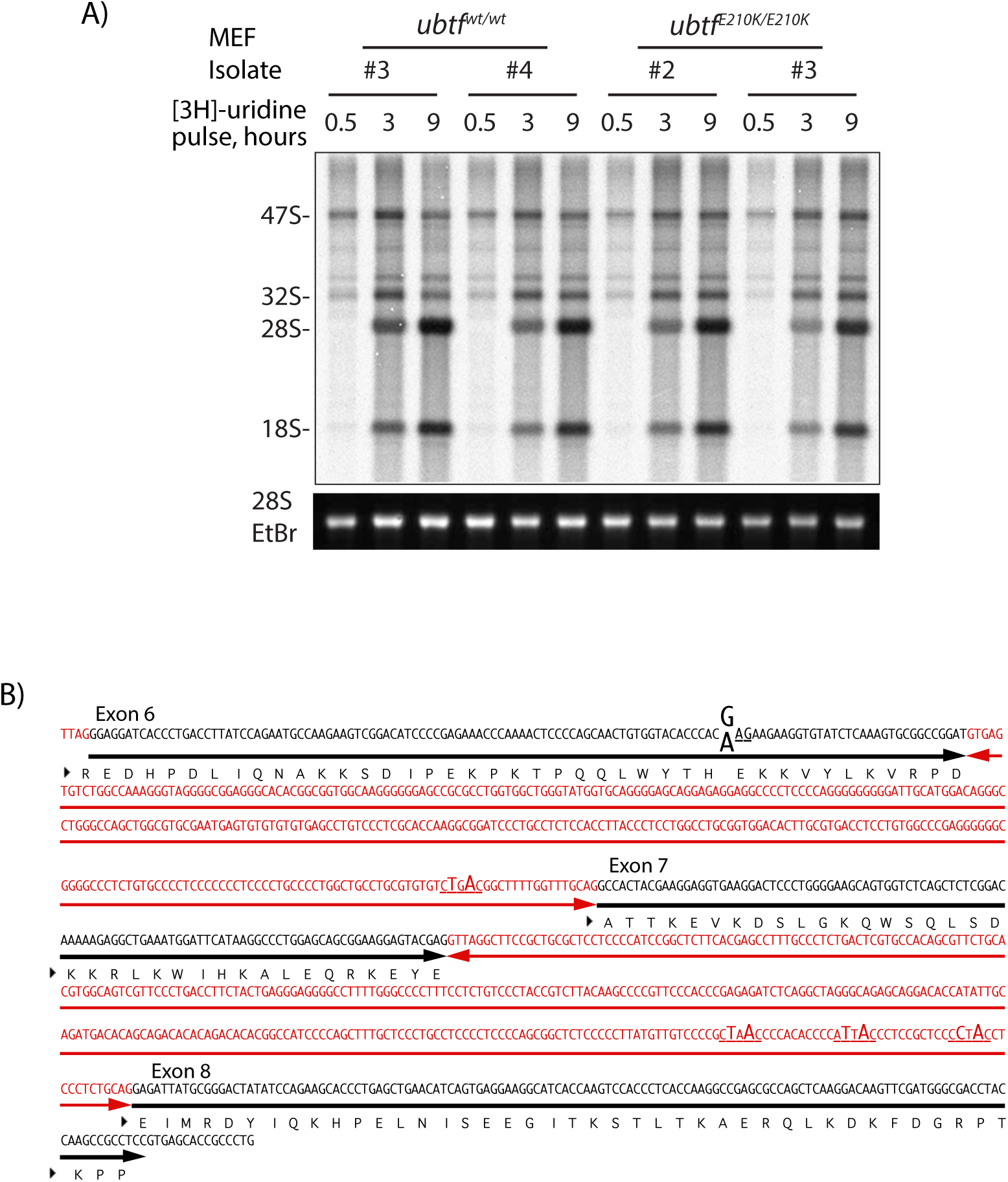
A) RNA metabolic pulse labelling to reveal 47S pre-rRNA synthesis and processing products in *Ubtf^E210k/E210K^* knock-in and wild type *Ubtf^wt/wt^* MEFs. Gel fractionation of RNA after increasing labelling times is shown for two individual (numbered) MEF isolates. B) DNA base sequence of the differentially spliced region of mouse *Ubtf* gene showing coding exon 6, the differentially spliced coding exon 7 and coding exon 8 in black and the intervening introns in red (taken from GRCm38:11:102303960:102320342). The position of the G>A gene mutation, the cause of the E210K change in the Ubtf protein, is indicated as are the potential splice branch sites in the intervening introns that most closely fit the yTnAy consensus (52).

**Figure S11.**
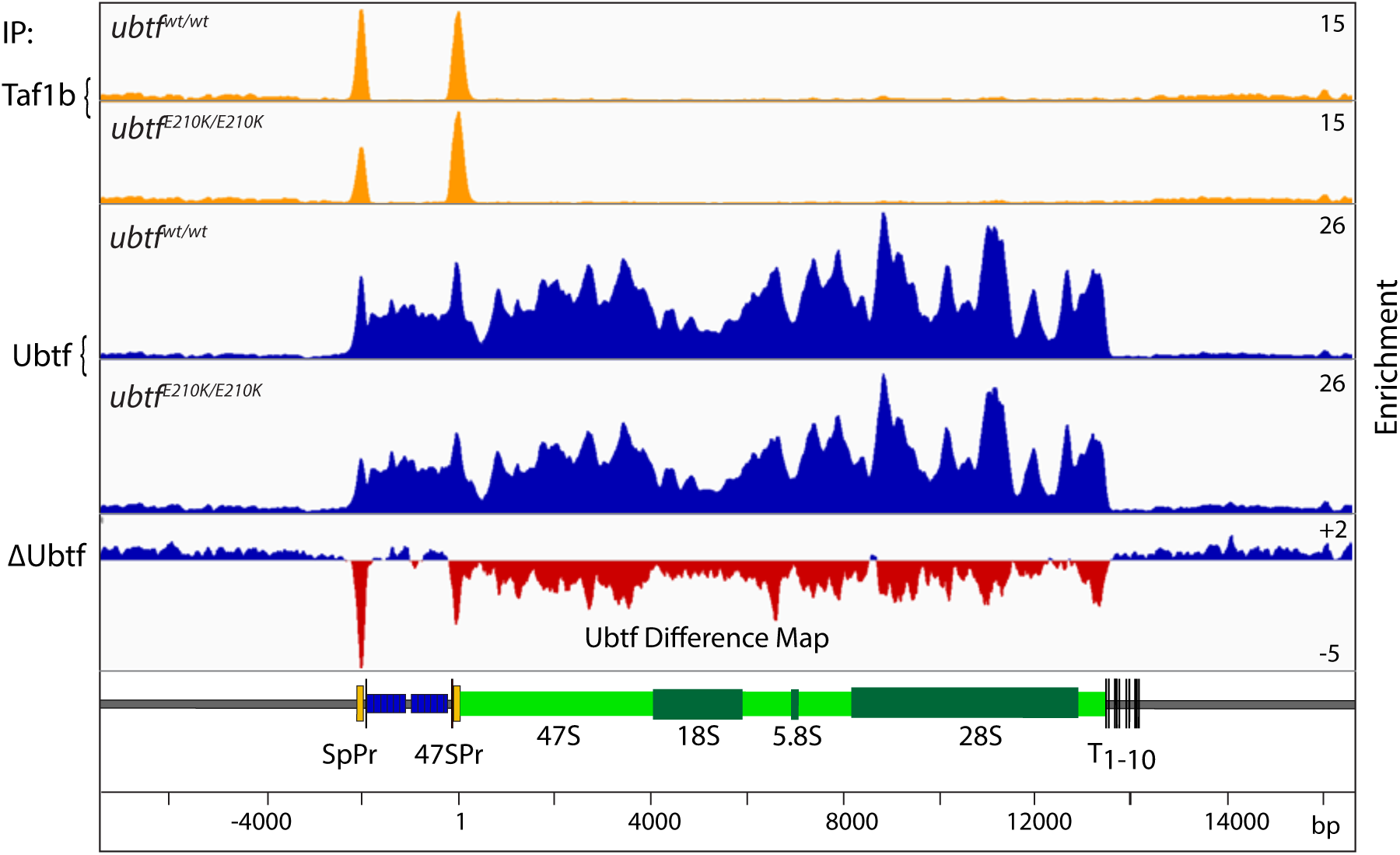
DChIP-Seq difference map of Ubtf occupancy (*Ubtf^E210K/E210K^* - wild type MEFs) as in Figure 6D but here shown across the full rDNA repeat. Taf1b and Ubtf mapping are also shown for reference.

## Notes

### Competing Interest Statement

The authors have declared no competing interest.

